## Additional_file_1 for "RNASeqR: an R package for automated two-group RNA-Seq analysis workflow"

RNASeqR : RNA-Seq analysis based on one independent variable


### RNASeqR : RNA-Seq analysis based on one independent variable

Author: Kuan-Hao Chao

###### *Last update: 30 November, 2018*

### Contents

- 1 Introduction
- 2 Sample Definition
- 3 System Requirements
- 4 Installation
- 5 “RNASeqRParam” S4 Object Creation
  - 5.1 `RNASeqRParam` Slots Explanation
  - 5.2 `RNASeqRParam` Constructor Checking
  - 5.3 Example
- 6 Environment Setup
  - 6.1 File Setup
  - 6.2 Installation of Necessary Tools
  - 6.3 Export Path
  - 6.4 Example
- 7 Quality Assessment of `FASTQ` sequence data
  - 7.1 “systemPipeR” Quality Assessment
  - 7.2 Example
- 8 Read Alignment and Quantification
  - 8.1 The *HISAT2* Indexer
  - 8.2 The *HISAT2* Aligner
  - 8.3 The *Rsamtools `SAM` to `BAM`* Converter
  - 8.4 The *StringTie* Assembler
  - 8.5 The *StringTie `GTF`* Merger
  - 8.6 The *Gffcompare* Comparer
  - 8.7 The *Ballgown input* Creator
  - 8.8 The *Reads Count Table* Creator
  - 8.9 Example
- 9 Gene-level Differential Analysis
  - 9.1 General Data Visualization
    - 9.1.1 Before Differential Expression Analysis
    - 9.1.2 After Differential Expression Analysis
  - 9.2 “ballgown” Analysis Based on FPKM Value
  - 9.3 “DESeq2” Analysis Based on Read Count
  - 9.4 “edgeR” Analysis Based on Read Count
  - 9.5 Example
- 10 Functional Analysis
  - 10.1 GO Enrichment Analysis
  - 10.2 KEGG Pathway Analysis
  - 10.3 Example
- 11 Conlusion
- 12 Session Information
- 13 References

### 1 Introduction

RNA-Seq is a revolutionary approach for discovering and investigating the transcriptome using next-generation sequencing technologies(Wang et al. 2009). Typically, this transcriptome analysis aims to identify genes differentially expressed across different conditions or tissues, resulting in an understanding of the important pathways that are associated with the conditions(Wang et al. 2009).

RNASeqR is an user-friendly R-based tool for running an RNA-Seq analysis pipeline including quality assessment, read alignment and quantification, differential expression analysis, and functional analysis. The main features of this package are an automated workflow, comprehensive reports with data visualization, and an extendable file structure. In this package, the new tuxedo pipeline published in Nature Protocols in 2016 can be fully implemented in the R environment, including extra functions such as reads quality assessment and functional analysis.

The following are the main tools that are used in this package: ‘HISAT2’ for read alignment(Kim et al. 2015); ‘StringTie’ for alignment assembly and transcripts quantification(Pertea et al. 2015); ‘Rsamtools’ for converting `SAM` files to `BAM` files(Morgan et al. 2018); ‘Gffcompare’ for comparing merged `GTF` files with reference `GTF` files; ‘systemPipeR’ for quality assessment(Backman et al. 2016); ‘ballgown’ (Fu et al. 2018), ‘DESeq2’ (Love et al. 2014) and ‘edgeR’ (Robinson et al. 2010;McCarthy et al. 2012) for finding potential differentially expressed genes; and ‘clusterProfiler’ (Yu et al. 2012) for Gene Ontology(GO) functional analysis and Kyoto Encyclopedia of Genes and Genomes(KEGG) pathway analysis.

The central concept behind this package is that **each step involved in RNA-Seq data analysis is a function call in R**. At the beginning, users will create a `RNASeqRParam` S4 object by running the `RNASeqRParam()` constructor function for all variable checking. After the creation of `RNASeqRParam`, it will be used as input to the following analysis functions.

1. `RNASeqEnvironmentSet_CMD()` or `RNASeqEnvironmentSet()`: to setup the RNA-Seq environment.
2. `RNASeqQualityAssessment_CMD()` or `RNASeqQualityAssessment()`: (optional) to run a quality assessment step.
3. `RNASeqReadProcess_CMD()` or `RNASeqReadProcess()`: to run reads alignment and quantification.
4. `RNASeqDifferentialAnalysis_CMD()` or `RNASeqDifferentialAnalysis()`: to run differential analysis via different R packages.
5. `RNASeqGoKegg_CMD()` or `RNASeqGoKegg()`: to conduct GO and KEGG analysis.

Functions with the `CMD` suffix create an R script and run `nohup R CMD BATCH script.R` in the background. Functions with no `CMD` suffix process in the R shell. After running the above functions, the whole RNA-Seq analysis is done and the files generated in each step are stored in an organized file directory. The `RNASeqR` package makes two-group comparative RNA-Seq analysis **more efficient** and **easier** for users.

Functions with `CMD` suffix will create an R script and run `nohup R CMD BATCH script.R` in background while functions with no `CMD` suffix will be processed in R shell. Files generated in each step will be kept in proper directory. Once the workflow is completed, a comprehensive RNA-Seq analysis is done. Additionally, this package is mainly designed for a two-group comparison setting, i.e. differential expression profile between two conditions.

### 2 Sample Definition

Sample data used in this vignette can be downloaded from the `RNASeqRData` experiment package. It originated from NCBI’s Sequence Read Archive for the entries SRR3396381, SRR3396382, SRR3396384, SRR3396385, SRR3396386, and SRR3396387. These samples are from Saccharomyces cerevisiae. Suitable reference genome and gene annotation files for this species can be further downloaded from iGenomes, Ensembl, R64-1-1. To create a mini dataset for demonstration purposes, reads aligned to the region from 0 to 100000 on chromosome XV were extracted. The analysis of this mini dataset will be shown in this vignette. The experimental data package is located here.

For more real case-control data and RNA-Seq analysis results from this package, please go to this website: https://github.com/HowardChao/RNASeqR\_analysis\_result.

### 3 System Requirements

Necessary:

1. R version >= 3.5.0
2. Operating System: Linux and macOS are supported in the `RNASeqR` package. Windows is not supported. (because StringTie and HISAT2 are not available for Windows).
3. Third-party software used in this package includes HISAT2, StringTie and Gffcompare. The availability of these commands will be checked by `system2()` through the R shell at the end of the ‘Environment Setup’ step. The environment must successfully built before running the following RNA-Seq analysis. By default, binaries will be installed based on the operating system of the workstation; therefore there is no additional compiling required. Alternatively, users can still decide to skip certain software binary installation. For more details, please refer to the ‘Environment Setup’ chapter.

Recommended:

1. Python: Python2 or Python3.
2. `2to3`: If the Python version on the workstation is 3, this command will be used. Generally, `2to3` is available if Python3 is available.
   - Python and `2to3` are used for creating raw read counts for DESeq2 and edgeR.
   - The following are two conditions that will create a raw read count from StringTie output.
     1. Python2
     2. Python3 with `2to3` command available.
   - If one of these conditions is met, the raw read count will be created and DESeq2 and edgeR will be run automatically in the ‘Gene-level Differential Analyses’ step. If not, DESeq2 and edgeR will be skipped during ‘Gene-level Differential Analysis’ step. Checking the Python version and `2to3` command on the workstation beforehand is highly recommended but not necessary.
3. HISAT2 indexex: Users are advised to provide an *‘indices/’* directory in *‘inputfiles/’*. HISAT2 requires at least 160 GB of RAM and several hours to index the entire human genome.

### 4 Installation

```
if (!require("BiocManager")) {
    install.packages("BiocManager")
}
BiocManager::install("RNASeqRData")
```

### 5 “RNASeqRParam” S4 Object Creation

This is the **first step** of the RNA-Seq analysis workﬂow in this package. Prior to conducting RNA-Seq analysis, it is necessary to implement a constructor function, called `RNASeqRParam()` and to create a `RNASeqRParam` S4 object which stores parameters not only for pre-checking but also for utilizing as input the parameters in the subsequent steps.

#### 5.1 `RNASeqRParam` Slots Explanation

There are 11 slots in `RNASeqRParam`:

1. os.type : The operating system type. Value is `linux` or `osx`. This package only supports ‘Linux’ and ‘macOS’ (not ‘Windows’). If another operating system is detected, ERROR will be reported.
2. `python.variable` : A Python-related variable. The value is a list of whether Python is available and the Python version (`TRUE` or `FALSE`, `2` or `3`).
3. `python.2to3` : Availability of the `2to3` command. THe value is `TRUE` or `FALSE`.
4. `path.prefix` : Path prefix of the *‘gene\_data/’*, *‘RNASeq\_bin/’*, *‘RNASeq\_results/’*, *‘Rscript/’* and *‘Rscript\_out/’* directories. It is recommended that you create a new, empty directory in which all the subsequent RNA-Seq results can be saved.

5. `input.path.prefix` : Path prefix of the *‘input\_files/’* directory. Users should prepare an *‘input\_file/’* directory with the following rules:
   - **`genome.name`.fa**: Reference genome in FASTA file formation.
   - **`genome.name`.gtf**: Gene annotation in GTF file formation.
   - **raw\_fastq.gz/**: Directory storing `FASTQ` files.

     - Supports paired-end read files only.
     - Names of paired-end `FASTQ` files : ’`sample.pattern`\_1.fastq.gz’ and ’`sample.pattern`\_2.fastq.gz’. `sample.pattern` must be distinct for each sample.
   - **phenodata.csv**: Information about the RNA-Seq experiment design.

     - First column : Distinct ids for each sample. The value of each sample of this column must match `sample.pattern` in `FASTQ` files in *‘raw\_fastq.gz/’*. Column names must be **ids**.
     - Second column : An independent variable for the RNA-Seq experiment. Values of each sample of this column can only be parameter `case.group` or `control.group`. The column name is the parameter `independent.variable`.
   - **indices/** : The directory storing `HT2` index files for the HISAT2 alignment tool.

     - This directory is **optional**. `HT2` index files corresponding to the reference genome can be installed at HISAT2 official website. Providing `HT2` files can accelerate the subsequent analysis steps. It is highly advised that you install `HT2` files.
     - If `HT2` index files are not provided, the *‘input\_files/indices/’* directory should be deleted.

6. `genome.name` : The genome name defined in this RNA-Seq workflow (ex. ‘`genome.name`.fa’, ‘`genome.name`.gtf’)
7. `sample.pattern` : A regular expression of paired-end fastq.gz files under *‘input\_files/raw\_fastq.gz/’*. **IMPORTANT!!** The expression shouldn’t have `_[1,2].fastq.gz` at the end.
8. `independent.variable`: Independent variable for the biological experiment design of a two-group RNA-Seq analysis workflow.
9. `case.group` : Name of the case group.
10. `control.group` : Name of the control group.
11. `indices.optional` : A logical value indicating whether *‘input\_files/indices/’* exists. Value is `TRUE` or `FALSE`

#### 5.2 `RNASeqRParam` Constructor Checking

1. Create a new directory for the RNA-Seq analysis. It is highly recommended to create a new, completely empty directory. The parameter `path.prefix` of `RNASeqRParam()` constructor should be the absolute path of this new directory. All the RNA-SeqR-related files are generated in the subsequent steps will be stored inside of this directory.
2. Create a valid *‘input\_files/’* directory. You should create a file directory named *‘input\_files/’* with the neccessary files inside. It should follow the rules mentioned above.
3. Run the constructor of the `RNASeqRParam` S4 object. This constructor will check the validity of the input parameters before creating the S4 objects.
   - Operating system
   - Python version
   - 2to3 command
   - Structure, content, and rules of *‘inputfiles/’*
   - Validity of input parameters

#### 5.3 Example

```
library(RNASeqR)
library(RNASeqRData)
```

```
input_files.path <- system.file("extdata/", package = "RNASeqRData")
rnaseq_result.path <- tempdir(check = TRUE)
list.files(input_files.path, recursive = TRUE)
```

```
##  [1] "input_files/Saccharomyces_cerevisiae_XV_Ensembl.fa" 
##  [2] "input_files/Saccharomyces_cerevisiae_XV_Ensembl.gtf"
##  [3] "input_files/phenodata.csv"                          
##  [4] "input_files/raw_fastq.gz/SRR3396381_XV_1.fastq.gz"  
##  [5] "input_files/raw_fastq.gz/SRR3396381_XV_2.fastq.gz"  
##  [6] "input_files/raw_fastq.gz/SRR3396382_XV_1.fastq.gz"  
##  [7] "input_files/raw_fastq.gz/SRR3396382_XV_2.fastq.gz"  
##  [8] "input_files/raw_fastq.gz/SRR3396384_XV_1.fastq.gz"  
##  [9] "input_files/raw_fastq.gz/SRR3396384_XV_2.fastq.gz"  
## [10] "input_files/raw_fastq.gz/SRR3396385_XV_1.fastq.gz"  
## [11] "input_files/raw_fastq.gz/SRR3396385_XV_2.fastq.gz"  
## [12] "input_files/raw_fastq.gz/SRR3396386_XV_1.fastq.gz"  
## [13] "input_files/raw_fastq.gz/SRR3396386_XV_2.fastq.gz"  
## [14] "input_files/raw_fastq.gz/SRR3396387_XV_1.fastq.gz"  
## [15] "input_files/raw_fastq.gz/SRR3396387_XV_2.fastq.gz"
```

Check the files in *‘inputfiles/’* directory.

```
exp <- RNASeqRParam(path.prefix = rnaseq_result.path, 
                    input.path.prefix = input_files.path, 
                    genome.name = "Saccharomyces_cerevisiae_XV_Ensembl", 
                    sample.pattern = "SRR[0-9]*_XV",
                    independent.variable = "state", 
                    case.group = "60mins_ID20_amphotericin_B", 
                    control.group = "60mins_ID20_control")
```

```
show(exp)
```

```
## RNASeqRParam S4 object
##               os.type : linux 
##       python.variable : (Availability: TRUE , Version: 3 )
##           python.2to3 : TRUE 
##           path.prefix : /tmp/RtmpY60M8j/ 
##     input.path.prefix : /home/kuan-hao/R/x86_64-pc-linux-gnu-library/3.5/RNASeqRData/extdata/ 
##           genome.name : Saccharomyces_cerevisiae_XV_Ensembl 
##        sample.pattern : SRR[0-9]*_XV 
##  independent.variable : state 
##            case.group : 60mins_ID20_amphotericin_B 
##         control.group : 60mins_ID20_control 
##      indices.optional : FALSE 
##  independent.variable : state
```

In this example, the `RNASeqRParam` S4 object is stored in `exp` for subsequent RNA-Seq analysis steps. Any ERROR occurring in the checking steps will terminate the program.

### 6 Environment Setup

This is the **second step** of the RNA-Seq analysis workflow in this package. To set up the environment, run `RNASeqEnvironmentSet_CMD()` to execute the process in the background, or run `RNASeqEnvironmentSet()` to execute the process in the R shell.

#### 6.1 File Setup

1. Create Base Directories *‘gene\_data/’*, *‘RNASeq\_bin/’*, *‘RNASeq\_results’*, *‘Rscript’*, and *‘Rscript\_out’* will be created in the `path.prefix` directory. Here is the usage of these five main directories:
   - *‘gene\_data/’*: Symbolic links of *‘input\_files/’* and files that are created in each step of RNA-Seq analysis will be stored in this directory.
   - *‘RNASeq\_bin/’*: The binaries of necessary tools, HISAT2, SAMtools, StringTie and Gffcompare, are installed in this directory.
   - *‘RNASeq\_results’*: The RNA-Seq results, for example, alignment results, quality assessment results, differential analysis results etc., will be stored in this directory.
   - *‘Rscript’*: If your run the `XXX_CMD()` function, the corresponding R script(`XXX.R`) for certain steps will be created in this directory.
   - *‘Rscript\_out’*: The corresponding output report for R scripts (`XXX.Rout`) will be stored in this directory.
2. Symbolic links will be created from files in *‘input\_files/’* to the `path.prefix` directory.

#### 6.2 Installation of Necessary Tools

The operating system of your workstation will be detected. If the operating system is not `Linux` or `macOS`, ERROR will be reported. Users can decide whether the installation of essential programs(HISAT2, StringTie and Gffcompare) is going be automatically processed.

Third-party softwares used in this package includes HISAT2, StringTie and Gffcompare. Binaries are available for these three programs, and by default, they will be installed automatically based on the operating system of the workstation. Zipped binaries will be unpacked and exported to the R environment PATH. **No compilation is needed**.

To specify, there are three parameters(`install.hisat2`, `install.stringtie` and `install.gffcompare`) in both the `RNASeqEnvironmentSet_CMD()` and `RNASeqEnvironmentSet()` functions for users to determine which software is going to be installed automatically or skipped. The default settings of these parameters are `TRUE` so that these three programs will be installed directly. Otherwise, users can skip certain software installation processes by turning the values to `FALSE`. Please make sure to check that the skipped programs are available using `system2()` through the R shell. Unavailability of any of the programs will cause failure in the ‘Environment Setup’ step.

Here is the version information of each software binary.

- HISAT2
  - Based on your operating system, `hisat2-2.1.0-Linux_x86_64.zip` or `hisat2-2.1.0-OSX_x86_64.zip` zipped file will be installed.
  - The downloaded file will be unzipped and all binary files will be copied to and run from *‘RNASeq\_bin/’*
- StringTie
  - Based on your operating system, `stringtie-1.3.4d.Linux_x86_64.tar.gz` or `stringtie-1.3.4d.Linux_x86_64` will be installed.
  - The downloaded file will be unzipped and all binary files will be copied to and run from *‘RNASeq\_bin/’*.
- Gffcompare
  - Based on your operating system, `gffcompare-0.10.4.Linux_x86_64.tar.gz` or `gffcompare-0.10.4.Linux_x86_64.tar.gz` will be installed.
  - The downloaded file will be unzipped and all binary files will be copied to and run from *‘RNASeq\_bin/’*.

#### 6.3 Export Path

*‘RNASeq\_bin/’* will be added to the R environment PATH so that these binaries can be found in the R environment in the R shell through `system2()`. In the last step of environment setup, the `hisat2 --version`,`stringtie --version`,`gffcompare --version`, and `samtools --version` commands will be checked in order to make sure the environment is correctly constructed. The environment must be set up successfully before the subsequent analyses.

#### 6.4 Example

Run `RNASeqEnvironmentSet_CMD()` or `RNASeqEnvironmentSet()`.

1. For runs in the background, the results will be reported in *‘Rscript\_out/Environment\_Set.Rout’*. Make sure the environment is successfully set up before running the subsequent steps.

```
RNASeqEnvironmentSet_CMD(exp)
```

2. For runs in the R shell, the results will be reported in the R shell. Make sure the environment is successfully setup before running the subsequent steps.

```
RNASeqEnvironmentSet(exp)
```

### 7 Quality Assessment of `FASTQ` sequence data

This is the **third step** of the RNA-Seq analysis workflow in this package. Different from other necessary steps, it is optional and can be run several times with each result stored separately. Although this step can be skipped, it is strongly recommended that it be performed before processing the alignment step. To evaluate the quality of the raw reads in the `FASTQ` files, run `RNASeqQualityAssessment_CMD()` in the background or run `RNASeqQualityAssessment()` to execute the process in the R shell.

#### 7.1 “systemPipeR” Quality Assessment

In this step, the systemPipeR package is used for evaluating sequencing reads and the details are as follows:

1. Check the number of times that the user has run the quality assessment process and create the corresponding files *‘RNASeq\_results/QA\_results/QA\_{times}’*.
2. RNA-Seq environment set up. The *‘rnaseq/’* directory will be created by systemPipeR package.
3. Create the *‘data.list.txt’* file.
4. Reading `FASTQ` files and create *‘fastqReport.pdf’* containing the results of the quality assessment.
5. Remove the *‘rnaseq/’* directory.

This quality assessment result (example below) is generated by systemPipeR package. It will be stored as a `PDF`.

#### 7.2 Example

Run `RNASeqQualityAssessment_CMD()` or `RNASeqQualityAssessment()`.

1. For runs in the background, the results will be reported in *‘Rscriptout/Quality\_Assessment.Rout’*. Make sure the quality assessment is successfully done before running the subsequent steps.

```
RNASeqQualityAssessment_CMD(exp)
```

2. Results fo runs in R shell will be reported in the R shell. Make sure the quality assessment is successfully done before running the subsequent steps.

```
RNASeqQualityAssessment(exp)
```

```
## [1] "Generated rnaseq directory. Next run in rnaseq directory, the R code from *.Rmd (*.Rnw) template interactively. Alternatively, workflows can be exectued with a single command as instructed in the vignette."
```

### 8 Read Alignment and Quantification

This is the **fourth step** of the RNA-Seq analysis workflow in this package. To process raw reads in `FASTQ` files, users can either run `RNASeqRawReadProcess_CMD()` to execute the process in background or run `RNASeqRawReadProcess()` to execute the process in the R shell. For further details about the commands and parameters that executed during each step, please check the reported *‘RNASeq\_results/COMMAND.txt’*.

#### 8.1 The *HISAT2* Indexer

In a preparation step (the `RNASeqRParam` creation step), the *‘indices/’* directory is checked for whether `HT2` index files already exist. If not, the following commands will be executed:

- Input: ‘`genome.name`.gtf’, ‘`genome.name`.fa’
- Output: ‘`genome.name`.ss’, ‘`genome.name`.exon’, ’`genome.name`\_tran.{number}.ht2’

1. `extract_splice_sites.py`, `extract_exons.py` execution
   - These two commands are executed to extract splice-site and exon information from the gene annotation file.
2. `hisat2-build` index creation
   - This command builds the HISAT2 index files *`genome.name`.ss* and *`genome.name`.exon* created in the previous step. Be aware that the index building step requires a larger amount of memory and longer time than other steps, and it might not be possible to run on some personal workstations. It is highly recommended that you check the availability of `HT2` index files at the HISAT2 official website for your target reference genome beforehands. Pre-installing `HT2` index files will greatly shorten the analysis time.
3. Write *‘RNASeq\_results/COMMAND.txt’*
   - Shell commands that run in this step will be documented in *‘RNASeq\_results/COMMAND.txt’*.

#### 8.2 The *HISAT2* Aligner

- Input: ’`genome.name`\_tran.{number}.ht2’, ‘`sample.pattern`.fastq.gz’
- Output: ‘`sample.pattern`.sam’

1. `hisat2` command is executed on paired end `FASTQ` files. `SAM` files will be created.
   - `SAM` files are stored in *‘gene\_data/raw\_bam/’*.
2. Write *‘RNASeq\_results/COMMAND.txt’*
   - The shell command that run in this step will be documented into *‘RNASeq\_results/COMMAND.txt’*.
3. The summary dataframe for alignment reads and mapping rates in terms of tabular(`CSV`) and picture (`PNG`) format is created and kept in the directory ‘RNASeq\_results/Alignment\_Report’.

#### 8.3 The *Rsamtools `SAM` to `BAM`* Converter

In this step, users can choose whether they want to use ‘Rsamtools’(an R package) or ‘SAMtools’ ( a command-line-based tool) to do files conversion by setting the `SAMtools.or.Rsamtools` parameter to `Rsamtools` or `SAMtools`. By default, `Rsamtools` will be used. However, if the amount of RNA-Seq data is too large, ‘Rsamtools’ might not be able to finish this process due to the Rtmp file issue, and therefore ‘SAMtools’ is recommended. Users have to make sure the ‘samtools’ command is available on the workstation beforehands or ERROR will be reported.

- Input: ‘`sample.pattern`.sam’
- Output: ‘`sample.pattern`.bam’

1. The Rsamtools package provides an interface to `samtools` in the R environment. In this step, `SAM` files from HISAT2 will be converted to `BAM` files by running the `asBam()` function.
   - Output `BAM` files are stored in *‘gene\_data/raw\_sam/’*.
2. Write *‘RNASeq\_results/COMMAND.txt’*

#### 8.4 The *StringTie* Assembler

- Input: ‘`genome.name`.gtf’, ‘`sample.pattern`.bam’
- Output: ‘`sample.pattern`.gtf’

1. `stringtie` command is executed.
   - The tool assembles RNA-Seq alignments into potential transcripts.
   - The assembled `GTF` files which are from each `FASTQ` files are stored in *‘gene\_data/raw\_gtf/’*
2. Write *‘RNASeq\_results/COMMAND.txt’*
   - Shell commands that run in this step will be documented in *‘RNASeq\_results/COMMAND.txt’*.

#### 8.5 The *StringTie `GTF`* Merger

- Input: ‘`sample.pattern`.gtf’
- Output: ‘stringtiemerged.gtf’, ‘mergelist.txt’

1. Create the file ‘mergelist.txt’.
   - *gene\_data/merged/mergelist.txt* is created for the merging step.
2. `stringtie` command is executed.
   - The transcript merger merges each *`sample.pattern`.gtf* into *stringtiemerged.gtf*
   - Output files are all stored in *‘gene\_data/merged/’*
3. Write *‘RNASeq\_results/COMMAND.txt’*
   - Shell commands that run in this step will be documented in *‘RNASeq\_results/COMMAND.txt’*.

#### 8.6 The *Gffcompare* Comparer

- Input: ‘`genome.name`.gtf’, ‘stringtie\_merged.gtf’
- Output: ‘merged.annotated.gtf’, ‘merged.loci’, ‘merged.stats’, ‘merged.stringtie\_merged.gtf.refmap’, ‘merged.stringtie\_merged.gtf.tmap’, ‘merged.tracking’

1. `gffcompare` command is executed.
   - The comparison result of the merged `GTF` file and reference annotation files is reported in the *‘merged/’* directory.
2. Write *‘RNASeq\_results/COMMAND.txt’*
   - Shell commands that run in this step will be documented in *‘RNASeq\_results/COMMAND.txt’*.

#### 8.7 The *Ballgown input* Creator

- Input: ‘stringtie\_merged.gtf’
- Output: ‘ballgown/’, ‘gene\_abundance/’

1. `stringtie` command is executed.
   - StringTie will create the input directory for the ballgown package for subsequent differential expression analysis. Ballgown-related files will be stored under each sample name in *‘gene\_data/ballgown/’*.
   - StringTie will store gene-related information in an `TSV` file under each sample name in *‘gene\_data/gene\_abundance/’*.
2. Write *‘RNASeq\_results/COMMAND.txt’*
   - Shell commands that run in this step will be documented in *‘RNASeq\_results/COMMAND.txt’*.

#### 8.8 The *Reads Count Table* Creator

Whether this step is executed depends on the availability of Python on your workstation.

- Input: ‘samplelst.txt’
- Output: ‘gene\_count\_matrix.csv’, ‘transcript\_count\_matrix.csv’

1. The reads count table converter Python script is downloaded as `prepDE.py`
2. Python checking
   - When Python is not available, this step is skipped.
   - When Python2 is available, `prepDE.py` is executed.
   - When Python3 is available, the `2to3` command will be checked.(Usually, if Python3 is installed, `2to3` command will be installed too.)
   - When Python3 is available but the `2to3` command is unavailable, the raw read count step will be skipped.
   - When Python3 and the `2to3` command are available, `prepDE.py` is converted to a file that can be executed by Python2 and is automatically executed.
3. Raw reads count creation
   - The raw reads count is created and the results are stored in *‘gene\_data/reads\_count\_matrix/’*
4. Write *‘RNASeq\_results/COMMAND.txt’*
   - Shell commands that run in this step will be documented in *‘RNASeq\_results/COMMAND.txt’*.

#### 8.9 Example

Run `RNASeqReadProcess_CMD()` or `RNASeqReadProcess()`.

1. For runs in the background, the results will be reported in *‘Rscriptout/Raw\_Read\_Process.Rout’*. Make sure the raw read process is successfully done before running the subsequent steps.

```
RNASeqReadProcess_CMD(exp)
```

2. For runs in the R shell, the results will be reported in R shell. Make sure the raw read process is successfully done before running the subsequent steps.

```
RNASeqReadProcess(exp)
```

### 9 Gene-level Differential Analysis

This is the **fifth step** of the RNA-Seq analysis workflow in this package. To identify differentially expressed genes, users can either run `RNASeqDifferentialAnalysis_CMD()` to execute the process in the background or run `RNASeqDifferentialAnalysis()` to execute the process in the R shell. In this package, we provide three normalized expression values: fragments per kilobase per million reads (FPKM) (Mortazavi et al. 2008), normalized counts by means of median of ratios normalization (MRN), or trimmed mean of m-values (TMM), each with proper statistical analyses using the R packages, ballgown, DESeq2, and edgeR. Gene IDs from StringTie and the ballgown R package will be mapped to ‘gene\_name’ in the GTF file for further functional analysis.

#### 9.1 General Data Visualization

Here we illustrate general data visualization before and after differential expression analysis. The results based on each differential analysis tool(*ballgown*, *DESeq2*, *edgeR*) are kept in the directory ‘RNASeq\_results/’. The plots shown below are the statistical results visualization of example data in the `RNASeqRData` package based on MRN-normalized expression values from DESeq2. For real data analysis results, please go to this website: https://howardchao.github.io/RNASeqR\_analysis\_result/.

##### 9.1.1 Before Differential Expression Analysis

###### 9.1.1.1 Frequency Plots

When visualizing the frequency of expression value per sample using the ggplot2 R package, the x-axis represents the range of normalized counts value by MRN or log_2_(MRN+1) value and the y-axis represents the frequency of each value on the x-axis

- **‘Frequency\_Plot\_normalized\_count\_ggplot2.png’**
- **‘Frequency\_Plot\_log\_normalized\_count\_ggplot2.png’**

###### 9.1.1.2 Distribution Plots

You can display the distribution of normalized expression values (e.g., log_2_(MRN+1) values) in a boxplot and a violin plot using the ggplot2 R package. Samples are colored as follows: blue for the case group and yellow for the control group.

- **‘Box\_Plot\_ggplot2.png’**
- **‘Violin\_Plot\_ggplot2.png’**

###### 9.1.1.3 PCA Plots

To display how the biological samples compare in overall similarities and differences using principal component analysis(PCA); the principal component scores of the top five dimensions are calculated using the FactoMineR package and the results are extracted and visualized using factoextra or ggplot2.

- **‘Dimension\_PCA\_Plot\_factoextra.png’**
- **‘PCA\_Plot\_factoextra.png’**: Samples are colored as follows: blue for the case group and yellow for the control group. The small point represents each sample while the big one represents each comparison group. Ellipases can be further added for grouping samples.
- **‘PCA\_Plot\_ggplot2.png’**: Samples are colored as described above.

###### 9.1.1.4 Correlation Plots

You can display Pearson correlation coefficient of a pairwise correlation analysis of changes in gene expression from all samples calculated by stats using ggplot2(correlation heat plot), corrplot(correlation dot plot), and PerformanceAnalytics(correlation bar plot). The colors from red to blue indicate the value of the correlation, from maximum value to minimum value, among all samples.

- **‘Correlation\_Heat\_Plot\_ggplot2.png’**
- **‘Correlation\_Dot\_Plot\_corrplot.png’**
- **‘Correlation\_Bar\_Plot\_PerformanceAnalytics.png’**

##### 9.1.2 After Differential Expression Analysis

###### 9.1.2.1 Volcano Plots

- **‘Volcano\_Plot\_graphics.png’**: You can display the criteria for selecting differentially expressed genes (DEGs) and identify these DEGs using ggplot2. The x-axis represents the log2-based fold change while the y-axis denotes the log10-based p-value. The up-regulated and down-regulated DEGs are hightlighted in red and green color, respectively.

###### 9.1.2.2 PCA Plots

To display how the biological samples compare in similarities and differences based on the expression value of DEGs using PCA. FactoMineR, factoextra and ggplot2 packages are used in this step.

- **‘Dimension\_PCA\_Plot\_factoextra.png’**
- **‘PCA\_Plot\_factoextra.png’**: Samples are colored as follows: blue for the case group and yellow for the control group. The small point represents each sample while the big one represents each comparison group. Ellipases can be further added for grouping samples.
- **‘PCA\_Plot\_ggplot2.png’**: Samples are colored as described above.

###### 9.1.2.3 Heatmap Plots

- **‘Heatmap\_Plot\_pheatmap.png’**: To visualize the expression value of DEGs in terms of the log2-based normalized value from all samples of the two groups using pheatmap. Samples are colored by defined groups: blue for the case group and yellow for the control group. Gene names and sample names will be shown on the heatmap except for two conditions: if there are more than 60 DEGs, gene names will not be shown, and if there are more than 16 samples, sample names will not be shown.

#### 9.2 “ballgown” Analysis Based on FPKM Value

Ballgown is an R package designed for differential expression analysis of RNA-Seq data. This package extracts FPKM values, i.e., read counts normalized by both library size and gene length, from StringTie software followed by applying a parametric F-test comparing the nested linear model as its default statistic model to identify DEGs. The basic steps are as follows:

- Input: *‘gene\_data/ballgown/’*
- Output: *‘RNASeq\_results/ballgown\_analysis/’*

1. Create a ballgown object that will be stored in *‘RNASeq\_results/ballgown\_analysis/ballgown.rda’*.
2. Filter out the genes for which the sum of FPKM values of all samples per gene equals 0.
3. Calculate log2-based fold change value in column `log2FC`.
4. Split a matrix of normalized counts into case and control groups based on phenotype data (*‘gene\_data/phenodata.csv’*) and assign relative information in column sample.pattern.FPKM.
5. Generate a `csv` file, *‘RNASeq\_results/ballgown\_analysis/ballgown\_normalized\_result.csv’*, in which to store normalized FPKM values, mean expression values per group, and statistical results.
6. Select DEGs based on default criteria: **`pval < 0.05` and `log2FC > 1 | log2FC < 1`**. Store the result in *‘RNASeq\_results/ballgown\_analysis/ballgown\_normalized\_DE\_result.csv’*
7. Additional data visualization: aside from the general data visualization mentioned above, transcript-related plots and an MA plot are also provided.

- **‘Distribution\_Transcript\_Count\_per\_Gene\_Plot.png’**: This plots the distribution of transcript counts per gene.
- **‘Distribution\_Transcript\_Length\_Plot.png’**: This plots the distribution of transcript length.
- **‘MA\_Plot\_ggplot2.png’**: This displays the difference of expression value between two groups by transforming the data onto log2-based ratio (x-axis) and log2-based mean (y-axis) scales by using ggplot2.

#### 9.3 “DESeq2” Analysis Based on Read Count

DESeq2 is an R package for count-based differential expression analysis using read counts to estimate variance-mean dependence. It takes sequence depth and gene composition into consideration and uses the MRN method to normalize read counts. The statistical model for differential expression is based on a negative binomial distribution. The basic steps are as follows:

- Input: *‘gene\_data/reads\_count\_matrix/gene\_count\_matrix.csv’*
- Output: *‘RNASeq\_results/ballgown\_analysis/’*

1. Create the `DESeqDataSet` object based on count data from the matrix of read counts and phenotype data using the `DESeqDataSetFromMatrix()` function.
2. Filter out the genes for which the sum of read counts of all samples equals 0.
3. Run `DESeq2()` function to process the differential expression analysis.
4. Generate a `csv` file, *‘RNASeq\_results/DESeq2\_analysis/DESeq2\_normalized\_result.csv’*, in which to store normalized MRN counts, mean expression values per group, and statistical results.
5. Select DEGs based on default criteria: **`pval < 0.05` and `log2FC > 1 | log2FC < 1`**. Store the result in *‘RNASeq\_results/DESeq2\_analysis/DESeq2\_normalized\_DE\_result.csv’*..
6. Additional data visualization: aside from the general data visualization mentioned above, a dispersion plot and an MA plot are also provided.

- **Dispersion\_Plot\_DESeq2.png**: This displays the dispersion estimates before and after normalization using `plotDispEsts()`. The x-axis denotes the mean of the normalized counts while the y-axis represents the dispersion estimates values using `plotDispEsts()` in DESeq2.
- **MA\_Plot\_DESeq2.png**: This displays the difference in expression values between two groups by transforming the data onto log2-based ratio (x-axis) and log2-based mean (y-axis) scales using `plotMA()` in DESeq2.

#### 9.4 “edgeR” Analysis Based on Read Count

edgeR is another R package for count-based differential expression analysis. It implements the TMM method to normalize count data between samples, and several statistical strategies based on the negative binomial distribution, such as exact tests, are used to detect differential expression. The basic steps are as follows:

- Input: *‘gene\_data/reads\_count\_matrix/gene\_count\_matrix.csv’*
- Output: *‘RNASeq\_results/edgeR\_analysis/’*

1. Create `DEGList` based on the count data from the matrix of read counts and phenotype data using the `DGEList()` function.
2. Normalize the `DEGList` object by running three functions in the following order: `calcNormFactors()`, `estimateCommonDisp()` and `estimateTagwiseDisp()`.
3. Conduct genewise exact tests using the `exactTest()` function.
4. Obtain a normalized count matrix using `cpm()` after TMM normalization. (cpm = counts per million)
5. Generate a `csv` file, ‘RNASeq\_results/edgeR\_analysis/edgeR\_normalized\_DE\_result.csv’, in which to store normalized TMM-normalized counts, mean expression values per group, and statistical results.
6. Select DEGs based on default criteria: **`pval < 0.05` and `log2FC > 1 | log2FC < 1`**. Store the result in ‘RNASeq\_results/edgeR\_analysis/edgeR\_normalized\_DE\_result.csv’.
7. Additional data visualization: aside from the general data visualization mentioned above, a MeanVar plot, a biological coefficient of variance (BCV) plot, an MDS plot, and a Smear plot are also provided.

- **MeanVar\_Plot\_edgeR.png**: This visualizes the mean-variance relationship before and after TMM normalization using the `plotMeanVar()` function in edgeR.
- **BCV\_Plot\_edgeR.png**: This displays the genewise biological coefficient of variance (BCV) against gene abundance using the `plotBCV()` function in edgeR.
- **MDS\_Plot\_edgeR.png**: This presents the expression differences between the samples using the `plotMDS.DGEList()` function in edgeR.
- **Smear\_Plot\_edgeR.png**: This plots the log2-based fold change against the log10-based concentration using `plotSmear()` in edgeR.

#### 9.5 Example

Run `RNASeqDifferentialAnalysis_CMD()` or `RNASeqDifferentialAnalysis()`.

1. For runs in the background, the results will be reported in *‘Rscriptout/Differential\_Analysis.Rout’*. Make sure the differential expression analysis is successfully finished before running subsequent steps.

```
RNASeqDifferentialAnalysis_CMD(exp)
```

2. For runs in the R shell, the results will be reported in the R shell. Make sure the differential expression analysis is successfully finished before running subsequent steps.

```
RNASeqDifferentialAnalysis(exp)
```

### 10 Functional Analysis

This is the **sixth step** of the RNA-Seq analysis workflow in this package. clusterProfiler is used for GO functional analysis and KEGG pathway analysis based on the DEGs found in three different differential analyses. Users can either run `RNASeqGoKegg_CMD()` to execute the process in the background or run `RNASeqGoKegg()` to execute the process in the R shell. In this step, users have to provide a gene name type, `input.TYPE.ID`, that is used in StringTie and ballgown and is supported in the `OrgDb.species` annotation package for the target species. In GO functional analysis and KEGG pathway analysis, the `input.TYPE.ID` ID type will be converted into an `ENTREZID` ID type by the `bitr()` function in clusterProfiler first. Those `input.TYPE.ID` with no corresponding `ENTREZID` will return `NA` and be filtered out. The genes with `Inf` or `-Inf` log2 fold change will be filtered out too. ID conversion is done in each differential analysis tools (ballgown, DESeq2 and edgeR).

In this example, the RNA-Seq analysis target species is *Saccharomyces cerevisiae*(yeast). The `OrgDb.species` is `org.Sc.sgd.db`; the `input.TYPE.ID` is `GENENAME`. IDs are converted from `GENENAME` to `ENTREZID`.

#### 10.1 GO Enrichment Analysis

GO is defined by three concepts relating to gene functions(GO terms) - molecular function(MF), cellular component(CC), and biological process(BP) - and by how these functions are related to each other. In this step, GO classiﬁcation and GO over-representation tests are conducted. To classify significant GO terms, DEGs are analyzed by the `groupGO()` function. Similarly, the GO over-representation test of DEGs is conducted using `enrichGO()`. Both results are stored in a `csv` file and the top 15 GO terms are visualized by both a bar chart and a bubble plot.

In this example, the DESeq2 CC GO Classification bar chart is shown.

In this example, the DESeq2 MF GO Over-representation bar chart and the DESeq2 MF GO Over-representation dot plot are shown.

#### 10.2 KEGG Pathway Analysis

KEGG is a database resource for understanding functions and utilities of a biological system from molecular-level information(KEGG website). In this step, KEGG over-representation tests can be conducted using the clusterProfiler package. A KEGG over-representation test of DEGs is conducted using `enrichKEGG()`. The result will be stored in a `csv` file. The pathway IDs that are found in the KEGG over-representation test will be visualized with the `pathview` package. KEGG pathway URL will also be stored.

In this example, due to the limited number of DEGs, no over-represented pathways are found.

#### 10.3 Example

Run `RNASeqGoKegg_CMD()` or `RNASeqGoKegg()`.

1. For runs in the background, the results will be reported in *‘Rscriptout/GO\_KEGG\_Analysis.Rout’*.

```
RNASeqGoKegg_CMD(exp, 
                 OrgDb.species = "org.Sc.sgd.db",
                 go.level = 3, 
                 input.TYPE.ID = "GENENAME",
                 KEGG.organism = "sce")
```

2. Results from runs in the R shell will be reported in R shell.

```
RNASeqGoKegg(exp, 
             OrgDb.species = "org.Sc.sgd.db", 
             go.level = 3, 
             input.TYPE.ID = "GENENAME",
             KEGG.organism = "sce")
```

### 11 Conlusion

RNASeqR is an user-friendly R-based tool for running case-control study(two group) RNA-Seq analysis pipelines. The six main steps in this package are *RNASeqRParam S4 Object Creation*, *Environment Setup*, *Quality Assessment*, *Reads Alignment and Quantification*, *Gene-level Differential Expression Analysis* and, *Functional Analysis.* The main features that RNASeqR provides are an automated workflow, an extendable file structure, comprehensive reports, and data visualization on widely-used differential analysis tools. With this R package, doing two-group RNA-Seq analysis will be much easier and faster.

### 12 Session Information

```
sessionInfo()
```

```
## R version 3.5.0 (2018-04-23)
## Platform: x86_64-pc-linux-gnu (64-bit)
## Running under: Ubuntu 17.10
## 
## Matrix products: default
## BLAS: /usr/lib/x86_64-linux-gnu/blas/libblas.so.3.7.1
## LAPACK: /home/kuan-hao/anaconda3/lib/libmkl_rt.so
## 
## locale:
##  [1] LC_CTYPE=en_US.UTF-8       LC_NUMERIC=C              
##  [3] LC_TIME=en_US.UTF-8        LC_COLLATE=C              
##  [5] LC_MONETARY=en_US.UTF-8    LC_MESSAGES=en_US.UTF-8   
##  [7] LC_PAPER=en_US.UTF-8       LC_NAME=C                 
##  [9] LC_ADDRESS=C               LC_TELEPHONE=C            
## [11] LC_MEASUREMENT=en_US.UTF-8 LC_IDENTIFICATION=C       
## 
## attached base packages:
##  [1] grid      parallel  stats4    stats     graphics  grDevices utils    
##  [8] datasets  methods   base     
## 
## other attached packages:
##  [1] DOSE_3.7.1           org.Sc.sgd.db_3.6.0  RNASeqRData_0.99.3  
##  [4] png_0.1-7            BiocStyle_2.9.6      RNASeqR_0.99.25     
##  [7] edgeR_3.23.3         limma_3.37.4         pathview_1.21.2     
## [10] org.Hs.eg.db_3.6.0   AnnotationDbi_1.43.1 IRanges_2.15.17     
## [13] S4Vectors_0.19.19    Biobase_2.41.2       BiocGenerics_0.27.1 
## [16] ggplot2_3.0.0       
## 
## loaded via a namespace (and not attached):
##   [1] reticulate_1.10             tidyselect_0.2.4           
##   [3] RSQLite_2.1.1               htmlwidgets_1.2            
##   [5] FactoMineR_1.41             BiocParallel_1.15.11       
##   [7] devtools_1.13.6             BatchJobs_1.7              
##   [9] munsell_0.5.0               units_0.6-0                
##  [11] systemPipeR_1.15.3          withr_2.1.2                
##  [13] colorspace_1.3-2            GOSemSim_2.7.1             
##  [15] Category_2.47.1             knitr_1.20                 
##  [17] rstudioapi_0.7              leaps_3.0                  
##  [19] labeling_0.3                KEGGgraph_1.41.0           
##  [21] urltools_1.7.1              GenomeInfoDbData_1.1.0     
##  [23] hwriter_1.3.2               bit64_0.9-7                
##  [25] pheatmap_1.0.10             rprojroot_1.3-2            
##  [27] xfun_0.3                    R6_2.2.2                   
##  [29] GenomeInfoDb_1.17.1         locfit_1.5-9.1             
##  [31] bitops_1.0-6                fgsea_1.7.1                
##  [33] gridGraphics_0.3-0          DelayedArray_0.7.39        
##  [35] assertthat_0.2.0            scales_1.0.0               
##  [37] ggraph_1.0.2                nnet_7.3-12                
##  [39] enrichplot_1.1.7            gtable_0.2.0               
##  [41] ballgown_2.13.1             sva_3.29.1                 
##  [43] systemPipeRdata_1.9.4       rlang_0.2.2                
##  [45] genefilter_1.63.2           BBmisc_1.11                
##  [47] scatterplot3d_0.3-41        splines_3.5.0              
##  [49] rtracklayer_1.41.5          lazyeval_0.2.1             
##  [51] acepack_1.4.1               europepmc_0.3              
##  [53] brew_1.0-6                  checkmate_1.8.5            
##  [55] BiocManager_1.30.2          yaml_2.2.0                 
##  [57] reshape2_1.4.3              GenomicFeatures_1.33.2     
##  [59] backports_1.1.2             rafalib_1.0.0              
##  [61] qvalue_2.13.0               Hmisc_4.1-1                
##  [63] clusterProfiler_3.9.2       RBGL_1.57.0                
##  [65] tools_3.5.0                 bookdown_0.7               
##  [67] ggplotify_0.0.3             RColorBrewer_1.1-2         
##  [69] ggridges_0.5.0              Rcpp_0.12.18               
##  [71] plyr_1.8.4                  base64enc_0.1-3            
##  [73] progress_1.2.0              zlibbioc_1.27.0            
##  [75] purrr_0.2.5                 RCurl_1.95-4.11            
##  [77] prettyunits_1.0.2           ggpubr_0.1.8               
##  [79] rpart_4.1-13                viridis_0.5.1              
##  [81] cowplot_0.9.3               zoo_1.8-3                  
##  [83] SummarizedExperiment_1.11.6 ggrepel_0.8.0              
##  [85] cluster_2.0.7-1             factoextra_1.0.5           
##  [87] magrittr_1.5                data.table_1.11.4          
##  [89] DO.db_2.9                   triebeard_0.3.0            
##  [91] matrixStats_0.54.0          evaluate_0.11              
##  [93] hms_0.4.2                   xtable_1.8-3               
##  [95] XML_3.98-1.16               gridExtra_2.3              
##  [97] compiler_3.5.0              biomaRt_2.37.6             
##  [99] tibble_1.4.2                crayon_1.3.4               
## [101] htmltools_0.3.6             GOstats_2.47.0             
## [103] mgcv_1.8-24                 Formula_1.2-3              
## [105] tidyr_0.8.1                 geneplotter_1.59.0         
## [107] sendmailR_1.2-1             DBI_1.0.0                  
## [109] tweenr_0.1.5                corrplot_0.85              
## [111] MASS_7.3-50                 PerformanceAnalytics_1.5.2 
## [113] ShortRead_1.39.0            Matrix_1.2-14              
## [115] quadprog_1.5-5              bindr_0.1.1                
## [117] igraph_1.2.2                GenomicRanges_1.33.13      
## [119] pkgconfig_2.0.2             flashClust_1.01-2          
## [121] rvcheck_0.1.0               GenomicAlignments_1.17.3   
## [123] foreign_0.8-71              xml2_1.2.0                 
## [125] roxygen2_6.1.0              annotate_1.59.1            
## [127] XVector_0.21.3              AnnotationForge_1.23.3     
## [129] stringr_1.3.1               digest_0.6.17              
## [131] graph_1.59.0                Biostrings_2.49.1          
## [133] rmarkdown_1.10              fastmatch_1.1-0            
## [135] htmlTable_1.12              GSEABase_1.43.1            
## [137] Rsamtools_1.33.5            commonmark_1.5             
## [139] rjson_0.2.20                nlme_3.1-137               
## [141] jsonlite_1.5                bindrcpp_0.2.2             
## [143] desc_1.2.0                  viridisLite_0.3.0          
## [145] pillar_1.3.0                lattice_0.20-35            
## [147] KEGGREST_1.21.1             httr_1.3.1                 
## [149] survival_2.42-6             GO.db_3.6.0                
## [151] glue_1.3.0                  xts_0.11-1                 
## [153] UpSetR_1.3.3                bit_1.1-14                 
## [155] Rgraphviz_2.25.1            ggforce_0.1.3              
## [157] stringi_1.2.4               blob_1.1.1                 
## [159] DESeq2_1.21.20              latticeExtra_0.6-28        
## [161] memoise_1.1.0               dplyr_0.7.6
```
