## Additional_file_2 for "RNASeqR: an R package for automated two-group RNA-Seq analysis workflow"

### Package ‘RNASeqR’

November 16, 2018

**Type** Package

**Title** RNASeqR: RNA-Seq workflow for case-control study

**Version** 1.0.0

**Date** 2018-8-7

**Author** Kuan-Hao Chao

**Maintainer** Kuan-Hao Chao <>

**biocViews** Genetics, Infrastructure, DataImport, Sequencing, RNASeq, GeneExpression, GeneSetEnrichment, Alignment, QualityControl, DifferentialExpression, FunctionalPrediction, ExperimentalDesign, GO, KEGG, Visualization, Normalization, Pathways, Clustering

**Description** This R package is designed for case-control RNA-Seq analysis (two-group).
There are six steps: ``RNASeqRParam S4 Object Creation'', ``Environment Setup'', ``Quality Assessment'',
``Reads Alignment & Quantification'', ``Gene-level Differential Analyses'' and ``Functional Analyses''.
Each step corresponds to a function in this package. After running functions in order,
a basic RNASeq analysis would be done easily.

**License** Artistic-2.0

**Encoding** UTF-8

**RoxygenNote** 6.1.0

**Depends** R(>= 3.5.0), ggplot2, pathview, edgeR, methods

**Imports** Rsamtools, tools, reticulate, ballgown, gridExtra, rafalib, FactoMineR, factoextra, corrplot, PerformanceAnalytics, reshape2, DESeq2, systemPipeR, systemPipeRdata, clusterProfiler, org.Hs.eg.db, org.Sc.sgd.db, stringr, pheatmap, grDevices, graphics, stats, utils, DOSE, Biostrings, parallel

**Suggests** knitr, png, grid, RNASeqRData

**VignetteBuilder** knitr

**SystemRequirements** RNASeqR only support Linux and macOS. Window is not supported. Python2 is highly recommended. If your machine is Python3, make sure '2to3' command is available.

**BugReports** https://github.com/HowardChao/RNASeqR/issues

**URL** https://github.com/HowardChao/RNASeqR

**NeedsCompilation** no

**git_url** https://git.bioconductor.org/packages/RNASeqR

**git_branch** RELEASE_3_8

**git_last_commit** 8df9801

**git_last_commit_date** 2018-10-30

**Date/Publication** 2018-11-15

### R topics documented:

**Index** **20**

---

`CheckToolAll` *CheckToolAll*

---

##### Description

Check whether 'Hisat2', 'Stringtie' and 'Gffcompare' are installed on the workstation

##### Usage

```
CheckToolAll(path.prefix, print = TRUE)
```

##### Arguments

| | |
|---|---|
| `path.prefix` | path prefix of 'gene_data/', 'RNASeq_bin/', 'RNASeq_results/', 'Rscript/' and 'Rscript_out/' directories. |
| `print` | If `TRUE`, detailed information will be printed. If `FALSE`, detailed information will not be printed. |

##### Value

None

##### Examples

```
data(yeast)
## Not run:
CheckToolAll(yeast@path.prefix,
             print=TRUE)
## End(Not run)
```

---

`RNASeqDifferentialAnalysis`
*RNASeqDifferentialAnalysis*

---

##### Description

This function will run differential analysis on ballgown, DESeq2 and edgeR in background.
This function do following things :

1. ballgown analysis
   Raw reads are normalized into FPKM values
   The main statistic test in ballgown is paramatic F-test comparing nested linear models

2. DESeq2 analysis
   Median of rations normalization(MRN) is used in DESeq2 for raw reads count normalization. Sequencing depth and RNA composition is taken into consideration is this normalization method.
   The main statistic test in DESeq2 is negative binomial distribution.

3. edgeR analysis
   Raw reads are normalized by TMM and library size. (run `calcNormFactors()` to get a DGEList, and then run `cpm()` on that DGEList)
   The main statistic test in edgeR is trimmed mean of M-values(TMM).

If you want to run differential analysis on ballgown, DESeq2, edgeR for the following RNA-Seq workflow in background, please see `RNASeqDifferentialAnalysis()` function.

##### Usage

```
RNASeqDifferentialAnalysis(RNASeqRParam, which.trigger = "OUTSIDE",
  INSIDE.path.prefix = NA, ballgown.run = TRUE, ballgown.pval = 0.05,
  ballgown.log2FC = 1, DESeq2.run = TRUE, DESeq2.pval = 0.1,
  DESeq2.log2FC = 1, edgeR.run = TRUE, edgeR.pval = 0.05,
  edgeR.log2FC = 1, check.s4.print = TRUE)
```

##### Arguments

`RNASeqRParam` S4 object instance of experiment-related parameters

`which.trigger` Default value is `OUTSIDE`. User should not change this value.

`INSIDE.path.prefix`
    Default value is `NA`. User should not change this value.

- `ballgown.run` Default `TRUE`. Logical value whether to run ballgown differential analysis.
- `ballgown.pval` Default `0.05`. Set the threshold of ballgown p-value to filter out differential expressed gene.
- `ballgown.log2FC` Default `1`. Set the threshold of ballgown log2 fold change to filter out differential expressed gene.
- `DESeq2.run` Default `TRUE`. Logical value whether to run DESeq2 differential analysis.
- `DESeq2.pval` Default `0.05`. Set the threshold of DESeq2 p-value to filter out differential expressed gene.
- `DESeq2.log2FC` Default `1`. Set the threshold of DESeq2 log2 fold change to filter out differential expressed gene.
- `edgeR.run` Default `TRUE`. Logical value whether to run edgeR differential analysis.
- `edgeR.pval` Default `0.05`. Set the threshold of edgeR p-value to filter out differential expressed gene.
- `edgeR.log2FC` Default `1`. Set the threshold of edgeR log2 fold change to filter out differential expressed gene.
- `check.s4.print` Default `TRUE`. If `TRUE`, the result of checking `RNASeqRParam` will be reported in 'Rscript_out/Environment_Set.Rout'. If `FALSE`, the result of checking `RNASeqRParam` will not be in 'Rscript_out/Environment_Set.Rout'.

#### Value

None

#### Author(s)

Kuan-Hao Chao

#### Examples

```
data(yeast)
## Not run:
RNASeqDifferentialAnalysis(RNASeqRParam = yeast)
## End(Not run)
```

---

### RNASeqDifferentialAnalysis_CMD

*RNASeqDifferentialAnalysis_CMD*

---

#### Description

This function will run differential analysis on ballgown, DESeq2 and edgeR in R shell.
This function do following things :

1. ballgown analysis
   Raw reads are normalized into FPKM values
   The main statistic test in ballgown is paramatic F-test comparing nested linear models

2. DESeq2 analysis
   Median of rations normalization(MRN) is used in DESeq2 for raw reads count normalization. Sequencing depth and RNA composition is taken into consideration is this normalization method.
   The main statistic test in DESeq2 is negative binomial distribution.

3. edgeR analysis
   Raw reads are normalized by TMM and library size. (run `calcNormFactors()` to get a DGEList, and then run `cpm()` on that DGEList)
   The main statistic test in edgeR is trimmed mean of M-values(TMM).

If you want to run differential analysis on ballgown, DESeq2, edgeR for the following RNA-Seq workflow in R shell, please see `RNASeqDifferentialAnalysis()` function.

##### Usage

```
RNASeqDifferentialAnalysis_CMD(RNASeqRParam, which.trigger = "OUTSIDE",
  INSIDE.path.prefix = NA, ballgown.run = TRUE, ballgown.pval = 0.05,
  ballgown.log2FC = 1, DESeq2.run = TRUE, DESeq2.pval = 0.1,
  DESeq2.log2FC = 1, edgeR.run = TRUE, edgeR.pval = 0.05,
  edgeR.log2FC = 1, run = TRUE, check.s4.print = TRUE)
```

##### Arguments

| | |
|---|---|
| `RNASeqRParam` | S4 object instance of experiment-related parameters |
| `which.trigger` | Default value is `OUTSIDE`. User should not change this value. |
| `INSIDE.path.prefix` | Default value is `NA`. User should not change this value. |
| `ballgown.run` | Default `TRUE`. Logical value whether to run ballgown differential analysis. |
| `ballgown.pval` | Default `0.05`. Set the threshold of ballgown p-value to filter out differential expressed gene. |
| `ballgown.log2FC` | Default `1`. Set the threshold of ballgown log2 fold change to filter out differential expressed gene. |
| `DESeq2.run` | Default `TRUE`. Logical value whether to run DESeq2 differential analysis. |
| `DESeq2.pval` | Default `0.05`. Set the threshold of DESeq2 p-value to filter out differential expressed gene. |
| `DESeq2.log2FC` | Default `1`. Set the threshold of DESeq2 log2 fold change to filter out differential expressed gene. |
| `edgeR.run` | Default `TRUE`. Logical value whether to run edgeR differential analysis. |
| `edgeR.pval` | Default `0.05`. Set the threshold of edgeR p-value to filter out differential expressed gene. |
| `edgeR.log2FC` | Default `1`. Set the threshold of edgeR log2 fold change to filter out differential expressed gene. |
| `run` | Default value is `TRUE`. If `TRUE`, 'Rscript/Environment_Set.R' will be created and executed. The output log will be stored in 'Rscript_out/Environment_Set.Rout'. If `False`, 'Rscript/Environment_Set.R' will be created without executed. |
| `check.s4.print` | Default `TRUE`. If `TRUE`, the result of checking `RNASeqRParam` will be reported in 'Rscript_out/Environment_Set.Rout'. If `FALSE`, the result of checking `RNASeqRParam` will not be in 'Rscript_out/Environment_Set.Rout'. |

#### Value

None

#### Author(s)

Kuan-Hao Chao

#### Examples

```
data(yeast)
## Not run:
RNASeqDifferentialAnalysis_CMD(RNASeqRParam = yeast)
## End(Not run)
```

---

RNASeqEnvironmentSet *RNASeqEnvironmentSet*

---

#### Description

Set up the environment for the following RNA-Seq workflow in R shell
This function do 4 things :

1. Create file directories.
2. Install necessary tools.
3. Export 'RNASeq_bin/' to the R environment.
4. Check command of tools.

First it will create 'gene_data/', 'RNASeq_bin/', 'RNASeq_results/', 'Rscript/', 'Rscript_out/' directories.
Afterwards, 'Hisat2', 'Stringtie', 'Gffcompare' will be installed under 'RNASeq_bin/Download/' and be unpacked under 'RNASeq_bin/Unpacked/'.
'RNASeq_bin/' will be added to the R environment and validity of tools will be checked.
Any ERROR occurs will be reported and the program will be terminated.
If you want to set up the environment for the following RNA-Seq workflow in background, please see `RNASeqEnvironmentSet_CMD()` function.

#### Usage

```
RNASeqEnvironmentSet(RNASeqRParam, which.trigger = "OUTSIDE",
  INSIDE.path.prefix = NA, install.hisat2 = TRUE,
  install.stringtie = TRUE, install.gffcompare = TRUE,
  check.s4.print = TRUE)
```

#### Arguments

| | |
|---|---|
| `RNASeqRParam` | S4 object instance of experiment-related parameters |
| `which.trigger` | Default value is `OUTSIDE`. User should not change this value. |
| `INSIDE.path.prefix` | Default value is `NA`. User should not change this value. |
| `install.hisat2` | Whether to install 'HISAT2' in this function step. Default value is`TRUE`. Set `FALSE` to skip 'HISAT2' installation. |
| `install.stringtie` | Whether to install 'StringTie' in this function step. Default value is `TRUE`. Set`FALSE` to skip 'StringTie' installation. |
| `install.gffcompare` | Whether to install 'Gffcompare' in this function step. Default value is `TRUE`. Set`FALSE` to skip 'Gffcompare' installation. |
| `check.s4.print` | Default `TRUE`. If `TRUE`, the result of checking `RNASeqRParam` will be reported in 'Rscript_out/Environment_Set.Rout'. If `FALSE`, the result of checking `RNASeqRParam` will not be in 'Rscript_out/Environment_Set.Rout'. |

#### Value

None

#### Author(s)

Kuan-Hao Chao

#### Examples

```
data(yeast)
## Not run:
RNASeqEnvironmentSet(RNASeqRParam = yeast)
## End(Not run)
```

---

`RNASeqEnvironmentSet_CMD`

*RNASeqEnvironmentSet_CMD*

---

#### Description

Set up the environment for the following RNA-Seq workflow in background.
This function do 4 things :

1. Create file directories.
2. Install necessary tools.
3. Export 'RNASeq_bin/' to the R environment.

4. Check command of tools.

First it will create 'gene_data/', 'RNASeq_bin/', 'RNASeq_results/', 'Rscript/', 'Rscript_out/' directories.
Afterwards, 'Hisat2', 'Stringtie', 'Gffcompare' will be installed under 'RNASeq_bin/Download/' and be unpacked under 'RNASeq_bin/Unpacked/'.
'RNASeq_bin/' will be added to the R environment and validity of tools will be checked.
Any ERROR occurs will be reported and the program will be terminated.
If you want to set up the environment for the following RNA-Seq workflow in R shell, please see `RNASeqEnvironmentSet()` function.

#### Usage

```
RNASeqEnvironmentSet_CMD(RNASeqRParam, install.hisat2 = TRUE,
  install.stringtie = TRUE, install.gffcompare = TRUE, run = TRUE,
  check.s4.print = TRUE)
```

#### Arguments

| | |
|---|---|
| `RNASeqRParam` | S4 object instance of experiment-related parameters |
| `install.hisat2` | Whether to install 'HISAT2' in this function step. Default value is`TRUE`. Set `FALSE` to skip 'HISAT2' installation. |
| `install.stringtie` | Whether to install 'StringTie' in this function step. Default value is `TRUE`. Set`FALSE` to skip 'StringTie' installation. |
| `install.gffcompare` | Whether to install 'Gffcompare' in this function step. Default value is `TRUE`. Set`FALSE` to skip 'Gffcompare' installation. |
| `run` | Default value is `TRUE`. If `TRUE`, 'Rscript/Environment_Set.R' will be created and executed. The output log will be stored in 'Rscript_out/Environment_Set.Rout'. If `False`, 'Rscript/Environment_Set.R' will be created without executed. |
| `check.s4.print` | Default `TRUE`. If `TRUE`, the result of checking `RNASeqRParam` will be reported in 'Rscript_out/Environment_Set.Rout'. If `FALSE`, the result of checking `RNASeqRParam` will not be in 'Rscript_out/Environment_Set.Rout'. |

#### Value

None

#### Author(s)

Kuan-Hao Chao

#### Examples

```
data(yeast)
## Not run:
RNASeqEnvironmentSet_CMD(yeast)
## End(Not run)
```

RNASeqGoKegg *RNASeqGoKegg*

##### Description

Run Gene Ontology(GO) and Kyoto Encyclopedia of Genes and Genomes(KEGG) analysis in R shell.
This function do Gene Ontology(GO) and Kyoto Encyclopedia of Genes and Genomes(KEGG) analysis :

1. Gene Ontology(GO) :
   (a) Do GO function classification analysis.
   (b) Do GO function enrichment analysis.
   (c) Visualization : bar plot, dot plot etc.
2. Kyoto Encyclopedia of Genes and Genomes(KEGG) :
   (a) Do KEGG pathway enrichment analysis
   (b) Pathway visulization with `pathview` package. KEGG webpage pathway url will also be created

If you want to do GO functional analysis and KEGG pathway analysis for the following RNA-Seq workflow in background, please see `RNASeqGoKegg_CMD()` function.

##### Usage

```
RNASeqGoKegg(RNASeqRParam, which.trigger = "OUTSIDE",
  INSIDE.path.prefix = NA, OrgDb.species, go.level = 3, input.TYPE.ID,
  KEGG.organism, check.s4.print = TRUE)
```

##### Arguments

`RNASeqRParam` S4 object instance of experiment-related parameters

`which.trigger` Default value is `OUTSIDE`. User should not change this value.

`INSIDE.path.prefix` Default value is `NA`. User should not change this value.

`OrgDb.species` the genome wide annotation packages of species on Bioconductor. Currently, there are 19 supported genome wide annotation packages of species.

`go.level` the depth of acyclic graph in GO analysis

`input.TYPE.ID` The gene name type in OrgDb.species annotation packahge.

`KEGG.organism` the species that are supported for KEGG analysis. Currently, there are more than 5000 supported species genome. Check the valid species terms on https://www.genome.jp/kegg/catalo

`check.s4.print` Default `TRUE`. If `TRUE`, the result of checking `RNASeqRParam` will be reported in 'Rscript_out/Environment_Set.Rout'. If `FALSE`, the result of checking `RNASeqRParam` will not be in 'Rscript_out/Environment_Set.Rout'

##### Value

None

##### Author(s)

Kuan-Hao Chao

##### Examples

```
data(yeast)
## Not run:
RNASeqGoKegg(RNASeqRParam  = yeast,
             OrgDb.species = "org.Sc.sgd.db",
             go.level = 3,
             input.TYPE.ID = "GENENAME",
             KEGG.organism = "sce")
## End(Not run)
```

---

`RNASeqGoKegg_CMD` *RNASeqGoKegg_CMD*

---

##### Description

Run Gene Ontology(GO) and Kyoto Encyclopedia of Genes and Genomes(KEGG) analysis in background.
This function do Gene Ontology(GO) and Kyoto Encyclopedia of Genes and Genomes(KEGG) analysis :

1. Gene Ontology(GO) :
   (a) Do GO function classification analysis.
   (b) Do GO function enrichment analysis.
   (c) Visualization : bar plot, dot plot etc.
2. Kyoto Encyclopedia of Genes and Genomes(KEGG) :
   (a) Do KEGG pathway enrichment analysis
   (b) Pathway visulization with `pathview` package. KEGG webpage pathway url will also be created

If you want to do GO functional analysis and KEGG pathway analysis for the following RNA-Seq workflow in R shell, please see `RNASeqGoKegg()` function.

##### Usage

```
RNASeqGoKegg_CMD(RNASeqRParam, OrgDb.species, go.level = 3,
  input.TYPE.ID, KEGG.organism, run = TRUE, check.s4.print = TRUE)
```

##### Arguments

| | |
|---|---|
| RNASeqRParam | S4 object instance of experiment-related parameters |
| OrgDb.species | the genome wide annotation packages of species on Bioconductor. Currently, there are 19 supported genome wide annotation packages of species. |
| go.level | the depth of acyclic graph in GO analysis |
| input.TYPE.ID | The gene name type in OrgDb.species annotation packahge. |
| KEGG.organism | the species that are supported for KEGG analysis. Currently, there are more than 5000 supported species genome. Check the valid species terms on https://www.genome.jp/kegg/catalo |
| run | Default value is TRUE. If TRUE, 'Rscript/Environment_Set.R' will be created and executed. The output log will be stored in 'Rscript_out/Environment_Set.Rout'. If False, 'Rscript/Environment_Set.R' will be created without executed. |
| check.s4.print | Default TRUE. If TRUE, the result of checking RNASeqRParam will be reported in 'Rscript_out/Environment_Set.Rout'. If FALSE, the result of checking RNASeqRParam will not be in 'Rscript_out/Environment_Set.Rout' |

##### Value

None

##### Author(s)

Kuan-Hao Chao

##### Examples

```
data(yeast)
## Not run:
RNASeqGoKegg_CMD(RNASeqRParam  = yeast,
                 OrgDb.species = "org.Sc.sgd.db",
                 go.level = 3,
                 input.TYPE.ID = "GENENAME",
                 KEGG.organism = "sce")

## End(Not run)
```

---

#### RNASeqQualityAssessment

*RNASeqQualityAssessment*

---

##### Description

Assess the quality of '.fastq.gz' files for RNA-Seq workflow in R shell. This step is optional in the whole RNA-Seq workflow.
This function reports the quality assessment result in packages systemPipeR For systemPipeR, 'RNASeq_results/QA_results/Rqc/systemPipeR/fastqReport.pdf' will be created.
If you want to assess the quality of '.fastq.gz' files for the following RNA-Seq workflow in background, please see RNASeqQualityAssessment_CMD() function.

##### Usage

```
RNASeqQualityAssessment(RNASeqRParam, which.trigger = "OUTSIDE",
  INSIDE.path.prefix = NA, check.s4.print = TRUE)
```

##### Arguments

| | |
|---|---|
| `RNASeqRParam` | S4 object instance of experiment-related parameters |
| `which.trigger` | Default value is `OUTSIDE`. User should not change this value. |
| `INSIDE.path.prefix` | Default value is `NA`. User should not change this value. |
| `check.s4.print` | Default `TRUE`. If `TRUE`, the result of checking `RNASeqRParam` will be reported in 'Rscript_out/Environment_Set.Rout'. If `FALSE`, the result of checking `RNASeqRParam` will not be in 'Rscript_out/Environment_Set.Rout' |

##### Value

None

##### Author(s)

Kuan-Hao Chao

##### Examples

```
data(yeast)
## Not run:
RNASeqQualityAssessment(RNASeqRParam = yeast)
## End(Not run)
```

---

#### RNASeqQualityAssessment_CMD

*RNASeqQualityAssessment_CMD*

---

##### Description

Assess the quality of '.fastq.gz' files for RNA-Seq workflow in background. This step is optional in the whole RNA-Seq workflow.
This function reports the quality assessment result in packages `systemPipeR` For `systemPipeR`, 'RNASeq_results/QA_results/Rqc/systemPipeR/fastqReport.pdf' will be created.
If you want to assess the quality of '.fastq.gz' files for the following RNA-Seq workflow in R shell, please see `RNASeqQualityAssessment()` function.

##### Usage

```
RNASeqQualityAssessment_CMD(RNASeqRParam, run = TRUE,
  check.s4.print = TRUE)
```

#### Arguments

| | |
|---|---|
| RNASeqRParam | S4 object instance of experiment-related parameters |
| run | Default value is TRUE. If TRUE, 'Rscript/Environment_Set.R' will be created and executed. The output log will be stored in 'Rscript_out/Environment_Set.Rout'. If False, 'Rscript/Environment_Set.R' will be created without executed. |
| check.s4.print | Default TRUE. If TRUE, the result of checking RNASeqRParam will be reported in 'Rscript_out/Environment_Set.Rout'. If FALSE, the result of checking RNASeqRParam will not be in 'Rscript_out/Environment_Set.Rout' |

#### Value

None

#### Author(s)

Kuan-Hao Chao

#### Examples

```
data(yeast)
## Not run:
RNASeqQualityAssessment_CMD(RNASeqRParam = yeast)
## End(Not run)
```

---

RNASeqR *RNASeqR-package*

---

#### Description

RNASeqR-package

---

RNASeqReadProcess *RNASeqReadProcess*

---

#### Description

Process raw reads for RNA-Seq workflow in R shell
This function do 5 things :

1. 'Hisat2' : aligns raw reads to reference genome. If indices.optional in RNASeqRParam is FALSE, Hisat2 indices will be created.
2. 'Rsamtools': converts '.sam' files to '.bam' files.
3. 'Stringtie': assembles alignments into transcript.

4. 'Gffcompare': examines how transcripts compare with the reference annotation.

5. 'Stringtie': creates input files for ballgown, edgeR and DESeq2.

6. raw reads count: create raw reads count for DESeq2 and edgeR

Before running this function, `RNASeqEnvironmentSet_CMD()` or `RNASeqEnvironmentSet()` must be executed successfully. If you want to process raw reads for the following RNA-Seq workflow in background, please see `RNASeqReadProcess_CMD()` function.

#### Usage

```
RNASeqReadProcess(RNASeqRParam, which.trigger = "OUTSIDE",
  INSIDE.path.prefix = NA, SAMtools.or.Rsamtools = "Rsamtools",
  num.parallel.threads = 1, Rsamtools.nCores = 1,
  Hisat2.Index.run = TRUE, Hisat2.Alignment.run = TRUE,
  Rsamtools.Bam.run = TRUE, StringTie.Assemble.run = TRUE,
  StringTie.Merge.Trans.run = TRUE, Gffcompare.Ref.Sample.run = TRUE,
  StringTie.Ballgown.run = TRUE, PreDECountTable.run = TRUE,
  check.s4.print = TRUE)
```

#### Arguments

`RNASeqRParam` S4 object instance of experiment-related parameters

`which.trigger` Default value is `OUTSIDE`. User should not change this value.

`INSIDE.path.prefix`
Default value is `NA`. User should not change this value.

`SAMtools.or.Rsamtools`
Default value is `Rsamtools`. User can set to `SAMtools` to use command-line-based 'samtools' instead.

`num.parallel.threads`
Specify the number of processing threads (CPUs) to use for each step. The default is 1.

`Rsamtools.nCores`
The number of cores to use when running 'Rsamtools' step.

`Hisat2.Index.run`
Whether to run 'HISAT2 index' step in this function step. Default value is `TRUE`. Set `FALSE` to skip 'HISAT2 index' step.

`Hisat2.Alignment.run`
Whether to run 'HISAT2 alignment' step in this function step. Default value is `TRUE`. Set `FALSE` to skip 'HISAT2 alignment' step.

`Rsamtools.Bam.run`
Whether to run 'Rsamtools SAM to BAM' step in this function step. Default value is `TRUE`. Set `FALSE` to skip 'Rsamtools SAM to BAM' step.

`StringTie.Assemble.run`
Whether to run 'StringTie assembly' step in this function step. Default value is `TRUE`. Set `FALSE` to skip 'StringTie assembly' step.

`StringTie.Merge.Trans.run`
Whether to run 'StringTie GTF merging' step in this function step. Default value is `TRUE`. Set `FALSE` to skip 'StringTie GTF merging' step.

`Gffcompare.Ref.Sample.run`
: Whether to run 'Gffcompare comparison' step in this function step. Default value is `TRUE`. Set `FALSE` to skip 'Gffcompare comparison' step.

`StringTie.Ballgown.run`
: Whether to run 'StringTie ballgown creation' step in this function step. Default value is `TRUE`. Set `FALSE` to skip 'StringTie ballgown creation' step.

`PreDECountTable.run`
: Whether to run 'gene raw reads count creation' step in this function step. Default value is `TRUE`. Set `FALSE` to skip 'gene raw reads count creation' step.

`check.s4.print`
: Default `TRUE`. If `TRUE`, the result of checking `RNASeqRParam` will be reported in 'Rscript_out/Environment_Set.Rout'. If `FALSE`, the result of checking `RNASeqRParam` will not be in 'Rscript_out/Environment_Set.Rout'.

#### Value

None

#### Author(s)

Kuan-Hao Chao

#### Examples

```
data(yeast)
## Not run:
## Before run this function, make sure \code{RNASeqEnvironmentSet_CMD()}
##(or\code{RNASeqEnvironmentSet()}) is executed successfully.
RNASeqReadProcess(RNASeqRParam         = yeast,
                  num.parallel.threads = 10)
## End(Not run)
```

---

`RNASeqReadProcess_CMD` *RNASeqReadProcess_CMD*

---

#### Description

Process raw reads for RNA-Seq workflow in background.
This function do 5 things :

1. 'Hisat2' : aligns raw reads to reference genome. If `indices.optional` in `RNASeqRParam` is `FALSE`, Hisat2 indices will be created.

2. 'Rsamtools': converts '.sam' files to '.bam' files.

3. 'Stringtie': assembles alignments into transcript.

4. 'Gffcompare': examines how transcripts compare with the reference annotation.

5. 'Stringtie': creates input files for ballgown, edgeR and DESeq2.

6. raw reads count: create raw reads count for DESeq2 and edgeR

Before running this function, `RNASeqEnvironmentSet_CMD()` or `RNASeqEnvironmentSet()` must be executed successfully.
If you want to process raw reads for the following RNA-Seq workflow in R shell, please see `RNASeqReadProcess()` function.

#### Usage

```
RNASeqReadProcess_CMD(RNASeqRParam, SAMtools.or.Rsamtools = "Rsamtools",
  num.parallel.threads = 1, Rsamtools.nCores = 1,
  Hisat2.Index.run = TRUE, Hisat2.Alignment.run = TRUE,
  Rsamtools.Bam.run = TRUE, StringTie.Assemble.run = TRUE,
  StringTie.Merge.Trans.run = TRUE, Gffcompare.Ref.Sample.run = TRUE,
  StringTie.Ballgown.run = TRUE, PreDECountTable.run = TRUE,
  run = TRUE, check.s4.print = TRUE)
```

#### Arguments

`RNASeqRParam` S4 object instance of experiment-related parameters

`SAMtools.or.Rsamtools`
Default value is `Rsamtools`. User can set to `SAMtools` to use command-line-based 'samtools' instead.

`num.parallel.threads`
Specify the number of processing threads (CPUs) to use for each step. The default is 1.

`Rsamtools.nCores`
The number of cores to use when running 'Rsamtools' step.

`Hisat2.Index.run`
Whether to run 'HISAT2 index' step in this function step. Default value is `TRUE`. Set `FALSE` to skip 'HISAT2 index' step.

`Hisat2.Alignment.run`
Whether to run 'HISAT2 alignment' step in this function step. Default value is `TRUE`. Set `FALSE` to skip 'HISAT2 alignment' step.

`Rsamtools.Bam.run`
Whether to run 'Rsamtools SAM to BAM' step in this function step. Default value is `TRUE`. Set `FALSE` to skip 'Rsamtools SAM to BAM' step.

`StringTie.Assemble.run`
Whether to run 'StringTie assembly' step in this function step. Default value is `TRUE`. Set `FALSE` to skip 'StringTie assembly' step.

`StringTie.Merge.Trans.run`
Whether to run 'StringTie GTF merging' step in this function step. Default value is `TRUE`. Set `FALSE` to skip 'StringTie GTF merging' step.

`Gffcompare.Ref.Sample.run`
Whether to run 'Gffcompare comparison' step in this function step. Default value is `TRUE`. Set `FALSE` to skip 'Gffcompare comparison' step.

`StringTie.Ballgown.run`
Whether to run 'StringTie ballgown creation' step in this function step. Default value is `TRUE`. Set `FALSE` to skip 'StringTie ballgown creation' step.

PreDECountTable.run
: Whether to run 'gene raw reads count creation' step in this function step. Default value is TRUE. Set FALSE to skip 'gene raw reads count creation' step.

run
: Default value is TRUE. If TRUE, 'Rscript/Environment_Set.R' will be created and executed. The output log will be stored in 'Rscript_out/Environment_Set.Rout'. If False, 'Rscript/Environment_Set.R' will be created without executed.

check.s4.print
: Default TRUE. If TRUE, the result of checking RNASeqRParam will be reported in 'Rscript_out/Environment_Set.Rout'. If FALSE, the result of checking RNASeqRParam will not be in 'Rscript_out/Environment_Set.Rout'.

#### Value

None

#### Author(s)

Kuan-Hao Chao

#### Examples

```
data(yeast)
## Not run:
## Before run this function, make sure \code{RNASeqEnvironmentSet_CMD()}
## (or\code{RNASeqEnvironmentSet()}) is executed successfully.
RNASeqReadProcess_CMD(RNASeqRParam = yeast,
                      num.parallel.threads = 10)
## End(Not run)
```

---

RNASeqRParam-class *RNASeqR*

---

#### Description

An S4 class for checking and storing RNA-Seq workflow parameters of this package.

#### Slots

- os.type 'linux' or 'osx'. The operating system type.
- python.variable A list storing python environment. (check.answer, python.version)
- python.2to3 Logical value whether 2to3 command is available on the workstation.
- path.prefix Path prefix of 'gene_data/', 'RNASeq_bin/', 'RNASeq_results/', 'Rscript/' and 'Rscript_out/' directories.
- input.path.prefix Path prefix of 'input_files/' directory,
- genome.name Variable of genome name defined in this RNA-Seq workflow (ex. genome.name.fa, genome.name.gtf).
- sample.pattern Regular expression of paired-end fastq.gz files under 'input_files/raw_fastq.gz'. Expression not includes _[1,2].fastq.gz.
- independent.variable Independent variable for the biological. experiment design of two-group RNA-Seq workflow.
- case.group Group name of the case group.
- control.group Group name of the control group.
- indices.optional Logical value whether 'indices/' is exit in 'input_files/'.

##### Author(s)

Kuan-Hao Chao

##### Examples

```
data(yeast)
"@"(yeast, os.type)
"@"(yeast, python.variable)
"@"(yeast, python.2to3)
"@"(yeast, path.prefix)
"@"(yeast, input.path.prefix)
"@"(yeast, genome.name)
"@"(yeast, sample.pattern)
"@"(yeast, independent.variable)
"@"(yeast, case.group)
"@"(yeast, control.group)
"@"(yeast, indices.optional)
```

---

#### RNASeqRParam-constructor

*RNASeqRParam*

---

##### Description

Constructor function for RNASeqRParam objects

##### Usage

```
RNASeqRParam(path.prefix = NA, input.path.prefix = NA,
  genome.name = NA, sample.pattern = NA, independent.variable = NA,
  case.group = NA, control.group = NA)
```

##### Arguments

| | |
|---|---|
| `path.prefix` | Path prefix of 'gene_data/', 'RNASeq_bin/', 'RNASeq_results/', 'Rscript/' and 'Rscript_out/' directories. |
| `input.path.prefix` | Path prefix of 'input_files/' directory. |
| `genome.name` | variable of genome name defined in this RNA-Seq workflow (ex. `genome.name`.fa, `genome.name`.gtf). |
| `sample.pattern` | Regular expression of paired-end fastq.gz files under 'input_files/raw_fastq.gz'. Expression not includes `_[1,2].fastq.gz`. |
| `independent.variable` | Independent variable for the biological experiment design of two-group RNA-Seq workflow. |
| `case.group` | Group name of the case group. |
| `control.group` | Group name of the control group. |

##### Value

an object of class `RNASeqRParam`

#### Author(s)

kuan-hao Chao

Kuan-Hao Chao

#### Examples

```
input_files.path <- system.file("extdata/", package = "RNASeqRData")
rnaseq_result.path <- tempdir(check = TRUE)
exp <- RNASeqRParam(path.prefix          = rnaseq_result.path,
                    input.path.prefix    = input_files.path,
                    genome.name          = "Saccharomyces_cerevisiae_XV_Ensembl",
                    sample.pattern       = "SRR[0-9]*_XV",
                    independent.variable = "state",
                    case.group           = "60mins_ID20_amphotericin_B",
                    control.group        = "60mins_ID20_control")
```

---

yeast *Toy RNASeqRParam object*

---

#### Description

Small RNASeqRParam S4 object created with checked valid parameters for demonstration purposes

#### Author(s)

Kuan-Hao Chao

#### Examples

```
data(yeast)
yeast
# RNASeqRParam S4 object for example demonstration.
```

### Index

CheckToolAll, 2

RNASeq (RNASeqRParam-class), 17
RNASeqDifferentialAnalysis, 3
RNASeqDifferentialAnalysis_CMD, 4
RNASeqEnvironmentSet, 6
RNASeqEnvironmentSet_CMD, 7
RNASeqGoKegg, 9
RNASeqGoKegg_CMD, 10
RNASeqQualityAssessment, 11
RNASeqQualityAssessment_CMD, 12
RNASeqR, 13
RNASeqReadProcess, 13
RNASeqReadProcess_CMD, 15
RNASeqRParam
(RNASeqRParam-constructor), 18
RNASeqRParam-class, 17
RNASeqRParam-constructor, 18

yeast, 19
