## Additional_file_3 for "RNASeqR: an R package for automated two-group RNA-Seq analysis workflow"

RNASeqRData : RNASeqR vignette sample data explanation


### RNASeqRData : RNASeqR vignette sample data explanation

Author: Kuan-Hao Chao

###### *Last update: 16 October, 2018*

### Contents

- 1 Introduction
- 2 `input_files` criteria
- 3 Sample definition
- 4 Sample data preparation process
- 5 Session Information

### 1 Introduction

RNASeqRData is a helper package for vignette in RNASeqR software package. This vignette shows the criteria of `input_files` and extraction process of mini example data.

### 2 `input_files` criteria

`input.path.prefix` is the parameter that stores the directory location of *‘input\_files/’*. Users have to prepare an *‘input\_file/’* before running RNASeqR package workflow. The criteria of *‘input\_file/’* are listed below:

- **`genome.name`.fa**: reference genome in FASTA file formation.
- **`genome.name`.gtf**: gene annotation in GTF file formation.
- **raw\_fastq.gz/**: directory storing `FASTQ` files.
  - Support paired-end reads files only.
  - Names of paired-end `FASTQ` files : ’`sample.pattern`\_1.fastq.gz’ and ’`sample.pattern`\_2.fastq.gz’. `sample.pattern` must be distinct for each sample.
- **phenodata.csv**: information about RNA-Seq experiment design.
  - First column : Distinct ids for each sample. Value of each sample of this column must match `sample.pattern` in `FASTQ` files in *‘raw\_fastq.gz/’*. Column names must be **ids**.
  - Second column : independent variable for the RNA-Seq experiment. Value of each sample of this column can only be parameter `case.group` and `control.group`. Column name is parameter `independent.variable`.
- **indices/** : directory storing `HT2` indices files for HISAT2 alignment tool.
  - This directory is **optional**. `HT2` indices files corresponding to target reference genome can be installed at HISAT2 official website. Providing `HT2` files can accelerate the subsequent steps. It is highly advised to install `HT2` files.
  - If `HT2` index files are not provided, *‘input\_files/indices/’* directory should be deleted.

### 3 Sample definition

The data in this experiment data package is originated from NCBI’s Sequence Read Archive for the entries SRR3396381, SRR3396382, SRR3396384, SRR3396385, SRR3396386, and SRR3396387. These samples were from *Saccharomyces cerevisiae*. To create mini data for demonstration purpose, reads aligned to the region from 0 to 100000 at chromosome XV were extracted. More details steps will be explained in the next chapter. Reference genome and gene annotation files, `Saccharomyces_cerevisiae_XV_Ensembl.fa` and `Saccharomyces_cerevisiae_XV_Ensembl.gtf`, are downloaded from iGenomes, Ensembl, R64-1-1.

### 4 Sample data preparation process

- `fastq.gz` files: `fastq.gz` files are aligned and analyzed in advanced in order to reduce the size of large raw fastq.gz, which are about 800M, and to keep the most differential expressed genes as far as possible. Reads aligned to the region from 0 to 100000 at chromosome XV were extracted only. Therefore, the size of these `fastq.gz` would be reduced to only about 5M. The following are the data processing steps:

1. SAMtools builds bam indexes of BAM files :

- ex: `samtools index SRR3396381.bam`

2. SAMtools extracts reads in certain range :

- ex: `samtools view -b SRR3396381.bam "XV:0-100000" > SRR3396381.extracted.bam`

3. SAMtools sorts extracted BAM files :

- ex: `samtools sort -n SRR3396381.extracted.bam -o SRR3396381.sorted.bam`

4. SAMtools gets splited fastq files :

- ex: `bedtools bamtofastq -i SRR3396381.sorted.bam -fq SRR3396381_XV_1.fastq -fq2 SRR3396381_XV_2.fastq`

5. gzip fastq files :

- ex: `gzip SRR3396381_XV_1.fastq`

Finally, mini data in this RNASeqRData package are created.

- `Saccharomyces_cerevisiae_XV_Ensembl.fa`: Only XV chromosome sequence are extracted.
- `Saccharomyces_cerevisiae_XV_Ensembl.gtf`: The whole *Saccharomyces cerevisiae* gtf annotation files.

### 5 Session Information

```
sessionInfo()
```

```
## R version 3.5.0 (2018-04-23)
## Platform: x86_64-pc-linux-gnu (64-bit)
## Running under: Ubuntu 17.10
## 
## Matrix products: default
## BLAS: /usr/lib/x86_64-linux-gnu/blas/libblas.so.3.7.1
## LAPACK: /usr/lib/x86_64-linux-gnu/lapack/liblapack.so.3.7.1
## 
## locale:
##  [1] LC_CTYPE=en_US.UTF-8       LC_NUMERIC=C              
##  [3] LC_TIME=en_US.UTF-8        LC_COLLATE=en_US.UTF-8    
##  [5] LC_MONETARY=en_US.UTF-8    LC_MESSAGES=en_US.UTF-8   
##  [7] LC_PAPER=en_US.UTF-8       LC_NAME=C                 
##  [9] LC_ADDRESS=C               LC_TELEPHONE=C            
## [11] LC_MEASUREMENT=en_US.UTF-8 LC_IDENTIFICATION=C       
## 
## attached base packages:
## [1] grid      stats     graphics  grDevices utils     datasets  methods  
## [8] base     
## 
## other attached packages:
## [1] png_0.1-7       BiocStyle_2.9.6
## 
## loaded via a namespace (and not attached):
##  [1] Rcpp_0.12.18       bookdown_0.7       digest_0.6.17     
##  [4] rprojroot_1.3-2    backports_1.1.2    magrittr_1.5      
##  [7] evaluate_0.11      stringi_1.2.4      rmarkdown_1.10    
## [10] tools_3.5.0        stringr_1.3.1      xfun_0.3          
## [13] yaml_2.2.0         compiler_3.5.0     BiocManager_1.30.2
## [16] htmltools_0.3.6    knitr_1.20
```
