## Additional_file_4 for "RNASeqR: an R package for automated two-group RNA-Seq analysis workflow"

### Package ‘RNASeqRData’

November 15, 2018

**Type** Package

**Title** RNASeqRData: sample data for RNASeqR software package demonstration

**Version** 1.0.0

**Author** Kuan-Hao Chao

**Maintainer** Kuan-Hao Chao <>

**biocViews** ExperimentData, Saccharomyces_cerevisiae_Data

**Description** RNASeqRData is a helper experiment package for vignette demonstration purpose in RNASeqR software package.

**Suggests** png, grid

**Depends** R (>= 3.5.0)

**License** Artistic-2.0

**Encoding** UTF-8

**NeedsCompilation** no

**RoxygenNote** 6.1.0

**git_url** https://git.bioconductor.org/packages/RNASeqRData

**git_branch** RELEASE_3_8

**git_last_commit** 34b47cc

**git_last_commit_date** 2018-10-30

**Date/Publication** 2018-11-15

#### R topics documented:
