## Supplementary figures and images for "RNASeqR: an R package for automated two-group RNA-Seq analysis workflow"

### Additional_file_5

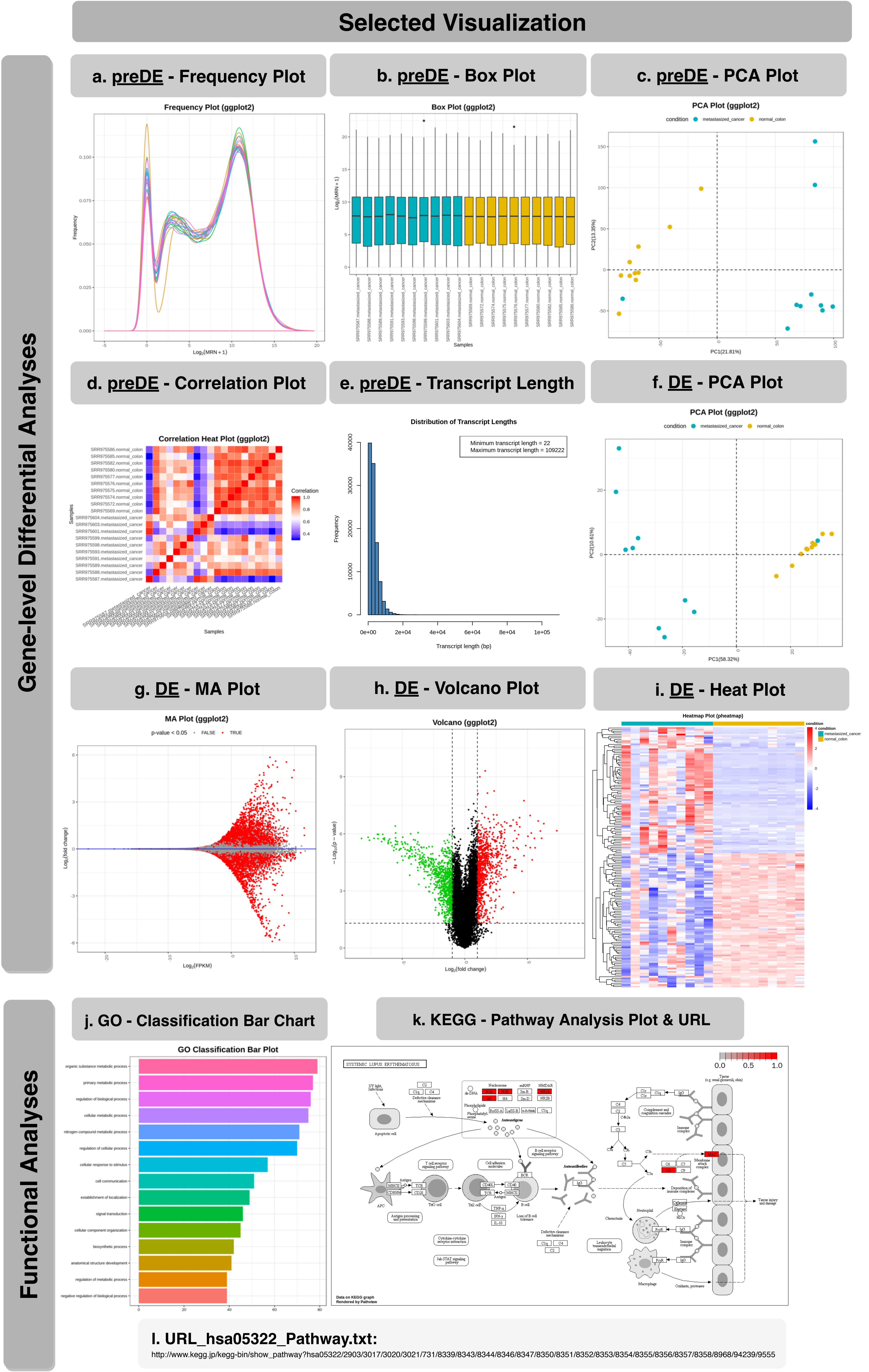

### Additional_file_6

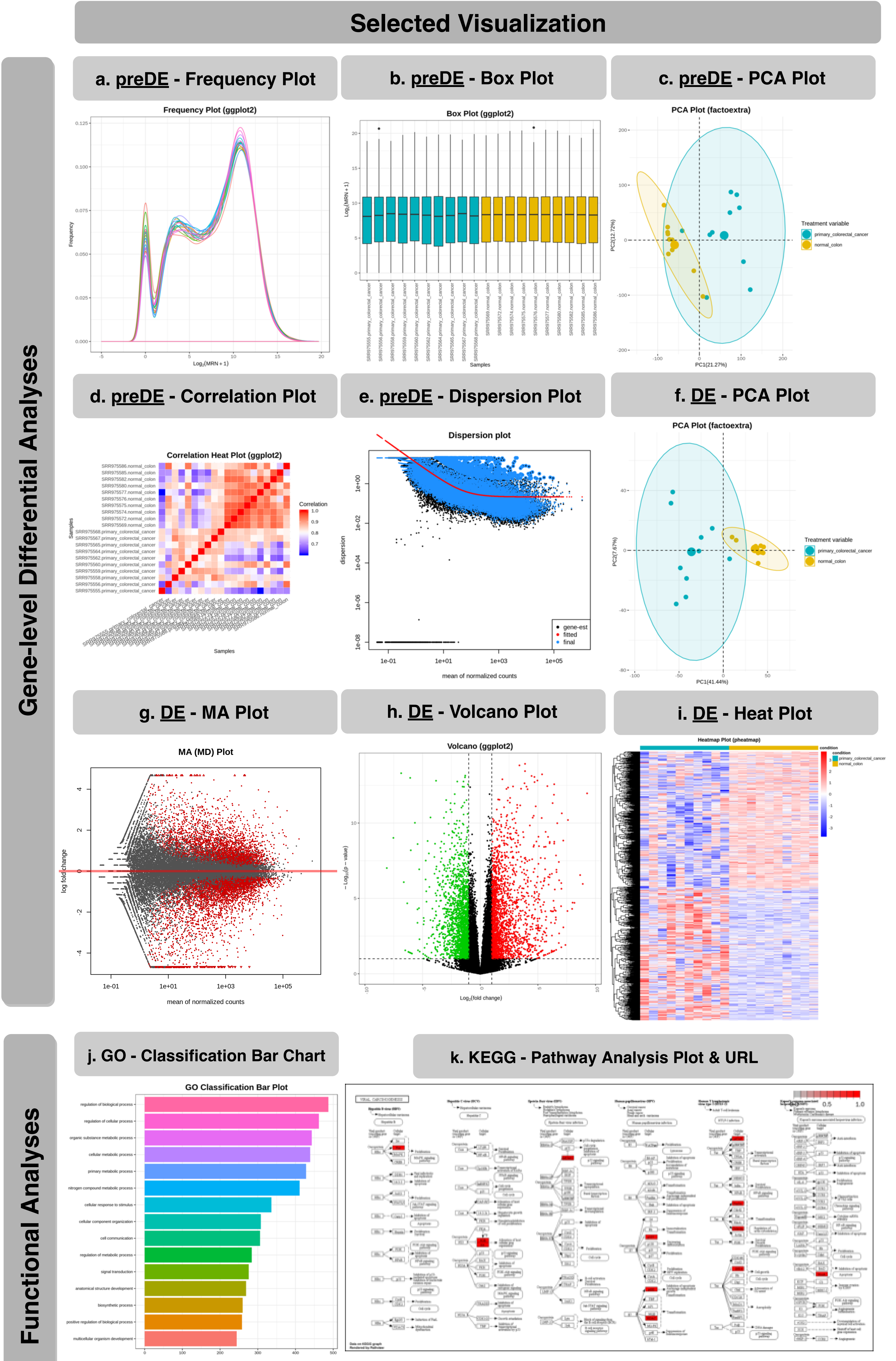


**l. URL_hsa05203_Pathway.txt:**

http://www.kegg.jp/kegg-bin/show_pathway?
hsa05203/1029/121504/23352/3017/3066/3845/4193/5366/554313/5922/5933/8294/8339/8343/8344/8346/8347/8359/8360/8361/8362/8363/8364/8365/8366/8367/8368/8370/85477/8850/9734

### Additional_file_7

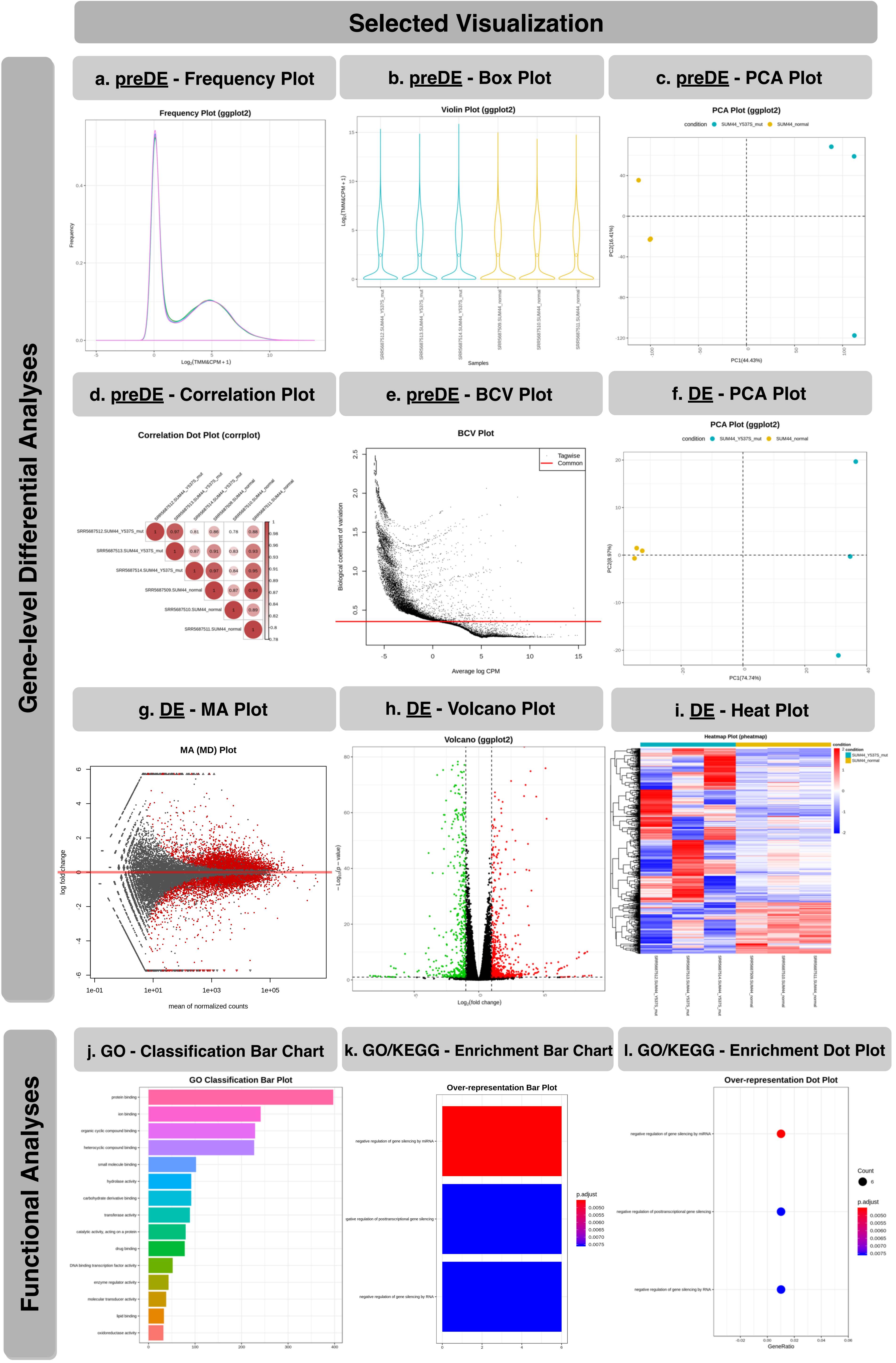
